# Defining Cardiac Recovery at Single Cell Resolution

**DOI:** 10.1101/2022.09.11.507463

**Authors:** Junedh M. Amrute, Lulu Lai, Pan Ma, Andrew L. Koenig, Kenji Kamimoto, Andrea Bredemeyer, Thirupura S. Shankar, Christoph Kuppe, Farid F. Kadyrov, Linda J. Schulte, Dylan Stoutenburg, Benjamin J. Kopecky, Sutip Navankasattusas, Joseph Visker, Samantha A. Morris, Rafael Kramann, Florian Leuschner, Douglas L. Mann, Stavros G. Drakos, Kory J. Lavine

**Author notes:** These authors contributed equally (Junedh M. Amrute, Lulu Lai). Correspondence should be addressed to: Kory J. Lavine, MD, PhD, Affiliation: Division of Cardiology, Department of Medicine, Washington University School of Medicine. Address: 660 S Euclid, Campus Box 8086, St. Louis, MO 63110.

## Abstract

Recovery of cardiac function is the ultimate goal of heart failure therapy. Unfortunately, cardiac recovery remains a rare and poorly understood phemomenon. Herein, we performed single nucleus RNA-sequencing (snRNA-seq) from non-diseased donors and heart failure patients. By comparing patients who recovered LV systolic function following LV assist device implantation to those who did not recover and donors, we defined the cellular and transcriptional landscape and predictors of cardiac recovery. We sequenced 40 hearts and recovered 185,881 nuclei with 13 distinct cell types. Using pseudobulk differential expression analysis to explicate cell specific signatures of cardiac recovery, we observed that recovered cardiomyocytes do not revert to a normal state, and instead, retain transcriptional signatures observed in heart failure. Macrophages and fibroblasts displayed the strongest signatures of recovery. While some evidence of reversion to a normal state was observed, many heart failure associated genes remained elevated and recovery signatures were predominately indicative of a biological state that was unique from donor and heart failure conditions. Acquisition of recovery states was associated with improved LV systolic function. Pro-inflammatory macrophages and inflammatory signaling in fibroblasts were identified as negative predictors of recovery. We identified downregulation of *RUNX1* transcriptional activity in macrophages and fibroblasts as a central event associated with and predictive of cardiac recovery. *In silico* perturbation of *RUNX1* in macrophages and fibroblasts recapitulated the transcriptional state of cardiac recovery. This prediction was corroborated in a mouse model of cardiac recovery mediated by BRD4 inhibition where we observed a decrease in macrophage and fibroblast *Runx1* expression, diminished chromatin accessibility within peaks linked to the *Runx1* locus, and acquisition of recovery signatures. These findings suggest that cardiac recovery is a unique biological state and identify RUNX1 as a possible therapeutic target to facilitate cardiac recovery.

## Introduction

Heart failure (HF) is a growing epidemic that affects over 23 million individuals worldwide and imparts a substantial burden on healthcare systems^1, 2^. Restoration of cardiac function, a phenomenon referred to as cardiac recovery, represents the holy grail for heart failure therapies. Cardiac recovery is characterized by restoration of normal (or near normal) left ventricular (LV) systolic function and reversal of LV dilatation, a process termed LV reverse remodeling. Cardiac recovery is infrequently observed following initiation of goal directed medical therapy in ambulatory patients and implantation of LV assist devices (LVADs) in patients with advanced heart failure^3–6^. Individuals who experience cardiac recovery have markedly improved survival and quality of life^7, 8^. At present much remains to be learned about cardiac recovery including the underlying cellular and molecular mechanisms, features that distinguish heart failure patients who will undergo recovery from those who will continue to experience disease progression, and whether cardiac recovery represents a reversion to normal or a unique biological state^9^. Unraveling the molecular basis of cardiac recovery in humans may help us identify new therapeutic targets for heart failure.

The advent of next generation sequencing has afforded researchers new opportunities to dissect the cellular diversity of human tissue across disease contexts^10, 11^. Recent studies have utilized single nucleus RNA sequencing (snRNA-seq) from healthy and failing human hearts to greatly expand our understanding of human heart failure and identified cell specific transcriptional signatures in genetic, dilated, and hypertrophic cardiomyopathies^12–16^. While some studies have explored mechanical unloading of the heart, there is a paucity of information regarding cardiac recovery at the single cell level. Designing and executing a study focused on cardiac recovery is challenging and must be carefully performed to discriminate cardiac recovery from mechanical unloading. To rigorously identify individuals who exhibit cardiac recovery, patients must undergo extensive phenotyping before and after LVAD implantation and paired tissue specimens from the same patient need to be collected from similar regions in the LV at the time of LVAD implantation and at the time of LVAD explant.

Herein, we defined the cellular and transcriptional landscape of cardiac recovery using snRNA-seq. We compared non-diseased donor controls with heart failure patients who experienced either cardiac recovery or ongoing heart failure before and after LVAD implantation (n=40). By performing cell specific pseudobulk differential expression analysis at the patient level, we uncovered that cardiac recovery represents a unique biological state and defined cell specific signatures of cardiac recovery that were distinct from both healthy and failing hearts. We further showed that transcriptional changes of cardiac recovery were encoded outside of the cardiomyocyte compartment predominately within macrophages and fibroblasts. We identified a *RUNX1* gene regulatory network (GRN) in macrophages and fibroblasts that was associated with and predictive of cardiac recovery using a deep neural network. *In silico* perturbation analysis^17^, revealed that *RUNX1* activity in macrophages and fibroblasts was predicted to control transition of cell states towards those associated with recovery and away from those associated with heart failure. Finally, we leveraged a mouse model of cardiac recovery^18^ to validate the putative role of the *Runx1* GRN in cardiac recovery. These findings uncover the biological state and cell specific mechanisms that contribute to cardiac recovery and identify *RUNX1* as a potential therapeutic target to facilitate cardiac recovery.

## Results

### Single nuclei RNA sequencing defines the cellular landscape of cardiac recovery

We perform snRNA-seq on paired transmural LV specimens obtained from the apical anterior wall of age- and sex-matched donor controls (n=14) and heart failure (HF) patients at the time of LVAD implant and at the time of LVAD explant. HF patients were divided into those who recovered LV systolic function (recovery/reverse remodeled, RR, n=5) and those demonstrated persistent reduction in LV ejection fraction (mechanically unloaded, U, n=8). Echocardiograms post-LVAD implant were performed using a predefined protocol where the LVAD speed was reduced to assess intrinsic LV systolic function^19, 20^ (Fig.1A). After doublet removal and quality control (Extended Data Fig. 1), we recovered 185,881 nuclei across 40 patient samples. We then performed dimensional reduction, UMAP construction, and cell clustering with differential gene expression to annotate cell types (Fig. 1B). We identified 13 transcriptionally distinct cell types marked by canonical gene markers (Fig. 1C, Extended Data Fig. 2A). We also constructed cell type specific gene set scores and detected strong separation across clusters (Extended Data Fig. 2B-C). Cell composition analysis showed a drop in cardiomyocytes and increase in the stromal cell fraction in HF pre- and post-LVAD samples relative to controls (Fig. 1D, Extended Data Fig. 2D). Echocardiographic data revealed the maked difference in LV ejection fraction post-LVAD implant between the recovered and unloaded groups (Fig. 1E). We then performed cell type specific pseudobulk differential gene expression analysis using the following comparisons: pre-LVAD HF (U-pre and RR-pre) versus donor, recovery post-LVAD (RR-post) versus pre-LVAD HF, and RR-post versus donor (Extended Data Fig. 3A-B).

**Figure 1.**
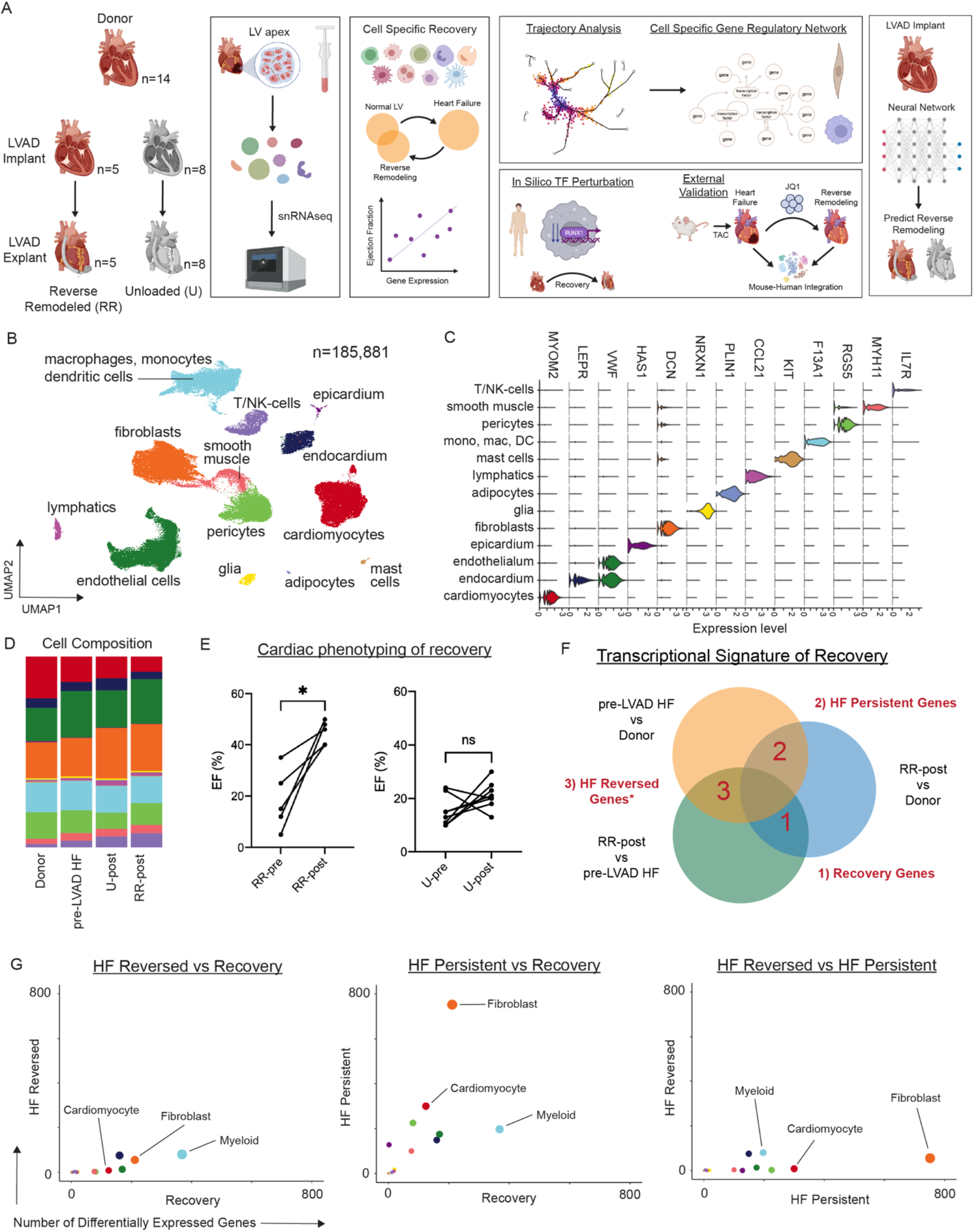
Study design, global clustering, and differential expression analysis of cardiac remodeling post-LVAD implantation. (A) Study design. (B) Integrated UMAP embedding plot of snRNA-seq data across n = 40 samples and 185,881 nuclei. (C) Violin plot for canonical marker genes for cell types. (D) Cell cluster composition across conditions colored by cell type (right). (E) Paired ejection fraction measured pre- and post-LVAD implantation split by RR (left) and U (right). Paired two-tailed t-test where *P = 0.0134 (RR) and nsP = 0.1169 (U). (F) Pseudobulk differential gene expression comparisons in cardiac remodeling post-LVAD implantation categories. (G) Number of statistically significant differentially expressed genes (p-adjusted value < 0.05 with and log2 fold-change > 0.58 from DEseq2) from (F) in each cell type as pairwise comparisons where size of dot refers to the sum of axis colored by cell type.

To place this DE analysis in the context of the different facets of cardiac recovery we built a Venn diagram of overlapping DE genes to identify three key categories relevant to the response to heart failure and cardiac recovery: (1) genes expressed specifically during recovery (recovery genes), (2) genes associated with HF that were persistently changed (HF persistent genes), and (3) genes associated with HF that returned to normal (HF reversed genes) (Fig. 1F). To identify cell types which may drive cardiac remodeling, we created pairwise scatter plots of the number of differentially expressed genes from the 3 comparisons (Fig. 1G). Within the majority of cell types, HF persistent and recovery genes were found at higher frequency than HF reversed genes indicating that the recovered state is not simply a reversion to normal and rather a unique biological entity.

### Cell type specific signatures of cardiac recovery

To dissect which cell types display the strongest signature of recovery, we quantified the number of recovery genes up- and down-regulated in each cell type. Among the major cell populations, myeloid cells (monocyte, macrophages, dendritic cells) and fibroblasts had the greatest number of recovery genes (Fig. 2A-B, Extended Data Fig. 3C). We then constructed heat maps of the top 25 up- and down-regulated recovery genes in the cells types with the strongest recovery signatures (myeloid, fibroblasts, endothelium, endocardium, cardiomyocytes) and observed consistent and robust enrichment within the RR-post group. We observed modest enrichment of the recovery signature within the U-post, suggesting that mechanical unloading may produce an intermediate phenotype within the continuum of recovery (Fig. 2C). To assess whether the transcriptional signature of recovery is unique to each cell type, we quantified the number of recovery genes which overlap in more than one cell type across the five major populations and found that most of the identified recovery genes are specific to a given cell type (Fig. 2D). Upset plots were constructed to assess pairwise overlap of cardiac recovery genes among different permutations of the major cell types (Extended Data Fig. 4). While the dominant signatures were cell type specific, we found 5 up-regulated (*PTPN13, FKBP5, ZBTB16, FGD4, ZMYND8*) and 1 down-regulated (*CTD-3252C9.4*) recovery genes common to the five major cell types (Fig. 2E). To ascertain whether cell type specific cardiac recovery signatures are associated with LV systolic function post-LVAD implant, we performed a linear regression analysis using LV ejection fraction as a surrogate for systolic function. We identified strong correlations between the ejection fraction and up- and down-regulated recovery signatures (Fig. 2F). These findings indicate that transcriptional signatures of cardiac recovery are cell type specific and are reflective of a continuum of cardiac recovery.

**Figure 2.**
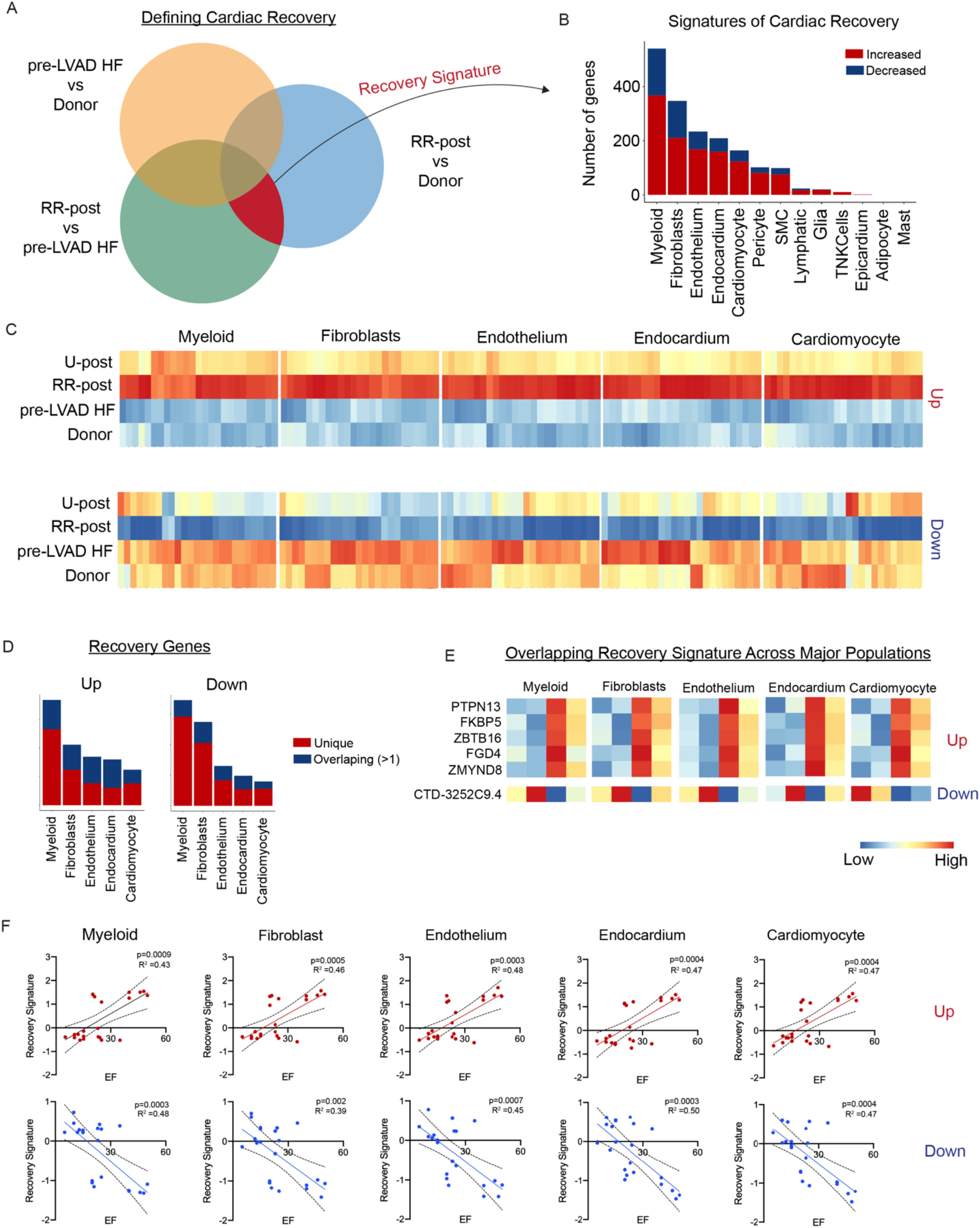
Cell type specific cardiac recovery. (A) Schematic of gene set from DE analysis which marks cardiac recovery. (B) Number of statistically significant (p-adjusted value < 0.05 with and log2 fold-change > 0.58 from DESeq2) genes from pseudobulk DE analysis which are up- and down-regulated in cardiac recovery across cell types. (C) Pseudobulk heatmaps of top genes up- (top) and down-regulated (bottom) in cardiac recovery in major cell populations split by donor, HF pre-LVAD, reverse remodeled, and unloaded. (D) Number of unique and overlapping cardiac recovery genes in major cell populations. (E) Pseudobulk expression of cardiac recovery genes which overlap among the major cell populations. (F) Polygenic recovery score of up- and down-regulated genes in cardiac recovery versus patient EF as a simple linear regression. Dotted line indicates 95% confidence interval, R^2^ goodness of fit, and p-value indicating whether the slope is significantly non-zero using an F test.

### Cardiomyocytes do not revert to a healthy state in cardiac recovery

To dissect the recovery landscape across cell populations, we performed a pseudobulk PCA at the sample level (Extended Data Fig. 5). We observed a strong separation between donor and HF samples across most cell types. Intriguingly, for cardiomyocytes, post-U and post-RR cardiomyocytes clustered with pre-LVAD HF cardiomyocytes suggesting a persistence of the HF phenotype during both recovery and mechanical unloading (Fig. 3A). To delineate whether recovery is associated with particular cardiomyocyte states, we performed high resolution clustering of cardiomyocytes. We detected 5 distinct cardiomyocyte cell states with unique transcriptional signatures and pathway enrichment (Fig. 3B-E). Gaussian kernel density plots showed only modest shifts in cell states between pre-LVAD HF, U-post, and RR-post conditions (Fig. 3F). We then created a pseudobulk heatmap of canonical genes up- and down-regulated in HF^13, 14^ and found that many HF genes are persistently dysregulated in recovered cardiomyocytes. Genes associated with the non-diseased donor state do not return to normal levels in recovery. Of note, reduced expression of NPPA and ANKRD1 was observed in pre- and post-LVAD implant in patients that recovered compared to those that did not recover, suggesting that the cardiomyocyte substrate may influence the propensity for recovery (Fig. 3G). We validated key findings using *in situ hybridization*. Compared to donor controls, the number of *MYH6* expressing cardiomyocytes was reduced in all pre- and post-LVAD HF samples. Consistent with our psudobulk analysis, the number of *NPPA* expressing cardiomyocytes was significantly increased in U-pre and U-post HF samples compared to donors. Modest trends were observed in RR-pre and RR-post samples (Fig. 3H-I). *MYH6* and NPPA expression were enriched in CM0 and CM1, respectively (Fig. 3J).

**Figure 3.**
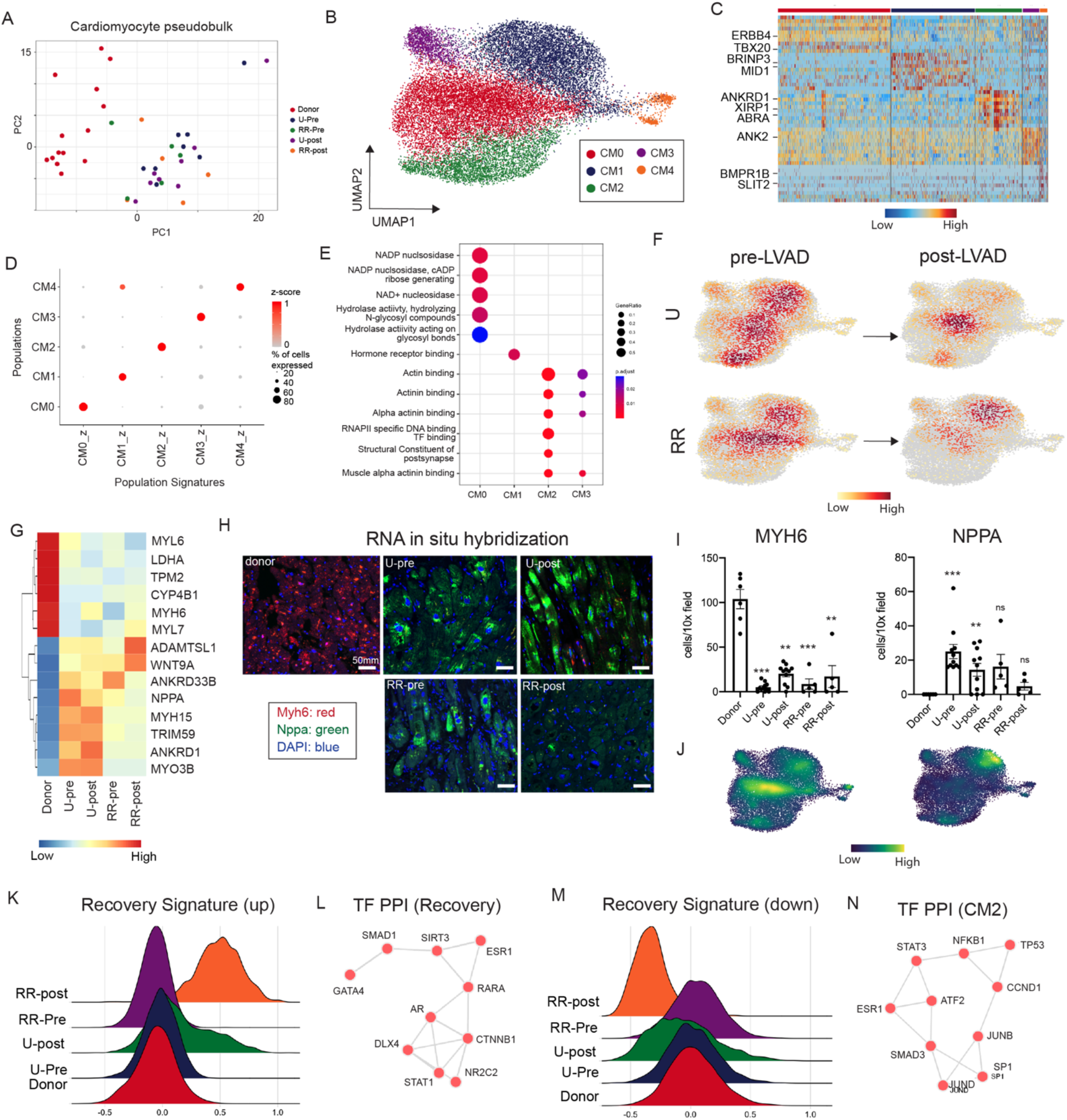
Cardiomyocytes do not revert to a healthy state in cardiac recovery. (A) Cardiomyocyte pseudobulk PCA representation of each patient sample colored by condition. (B) UMAP embedding plot of cardiomyocytes. (C) Heatmap of marker genes for distinct cardiomyocyte cell states. (D) Dot plot of gene set z-scores for top 10 genes in each cardiomyocyte cell state. (E) Enriched GO pathways for cardiomyocyte cell states using statistically significant marker genes identified using a Wilcoxon Rank Sum test (adjusted p-value < 0.05 and log2 fold-change > 0.58). Dot size refers to gene ratio and color of dots refers to the adjusted p-value. (F) Gaussian kernel density estimation of number of nuclei in the UMAP embedding split by condition. (G) Pseudobulk heatmap of canonical genes up and down in heart failure split by condition. (H) Fluorescent RNAscope in situ hybridization for MYH6 and NPPA in donor, pre-LVAD unloaded, pre-LVAD reverse remodeled, post-LVAD unloaded and post-LVAD reverse remodeled. Images are at 10x magnification. (I) Quantification of number cells per 10x field for MYH6 and NPPA across the five conditions. Brown-Forsythe and Welch ANOVA test comparing each condition to donor. Asterisks indicate degree of statistical significance. For MYH6 n = 38 total samples and Brown-Forsythe ANOVA test F = 28.91, DFn = 4, and p < 0.0001; U-pre (***P = 0.001), U-post (**P = 0.0011), RR-pre (***P = 0.0004), and RR-post (**P = 0.0019) relative to donor. For NPPA n = 38 total samples and Brown-Forsythe ANOVA test F = 6.040, DFn = 4, and p = 0.004; U-pre (***P = 0.0001), U-post (**P = 0.0033), RR-pre (ns P = 0.086), and RR-post (ns P = 0.1017) relative to donor. (J) Density plots of MYH6 and NPPA expression in UMAP embedding. (K) Up regulated pseudobulk recovery signature Ridge plot split across five conditions. (L) Transcription factor protein-protein interactions for genes up regulated in recovery. (M) Down regulated pseudobulk recovery signature Ridge plot split across five conditions. (N) Transcription factor protein-protein interactions for CM2 marker genes downregulated in cardiac recovery.

Using the up- and down-regulated genes in recovery from our pseudobulk analysis, we created a recovery signature for cardiomyocytes that demonstrated selective enrichment in RR-post group (Fig. 3K, M). To examine transcription factors which may modulate recovery, we used EnrichR to find transcription factors predicted to regulate genes up- and down-regulated in recovery. Notably, we observed up-regulation of a *GATA4* associated transcriptional network (Fig. 3L) and down-regulation of transcription factors associated with inflammation (*STAT3*, *JUNB*, *JUND*, *NFKB1*) in recovery (Fig. 3N). The gene signature downregulated in recovery was most enriched in the CM2 subset, which expressed canonical HF genes including *ANKRD1* and *NPPA* (Extended Data Fig. 6A). Among the genes down-regulated in cardiac recovery, *ABRA* was also expressed in CM2 and found to be reduced in HF patients that recovered pre- and post-LVAD implant (Extended Data Fig. 6B-C). To validate that reduced cardiomyocyte *ABRA* expression is predictive of and associated with cardiac recovery, we performed *in situ hybridization*. Quantification of *ABRA* expressing cardiomyocytes confirmed reductions in RR-pre and RR-post compared to U-pre and U-post groups (Extended Data Fig. 6D-E).

### Macrophages encode signatures that associate with and predict cardiac recovery

Pseudobulk PCA identified that RR-post myeloid cells clustered independent from donor or pre-LVAD HF myeloid cells, indicating a strong transcriptional signature of recovery in the myeloid compartment. U-post myeloid cells clustered with both RR-post and pre-LVAD myeloid cells consistent with the finding that recovery transcriptional signatures identify a spectrum of recovery (Fig. 4A). To decipher elements of cardiac recovery encoded in the monocyte, macrophage, and dendritic cell compartment, we sub-clustered and identified 9 distinct cell states with unique transcriptional signatures and pathway enrichment (Fig. 4B-C, Extended Data Fig. 7A-B). Cell composition analysis showed expansion of Mac1 and Mac2 and reduction of Mac5 in the U-pre and U-post groups relative to donors, RR-pre, and RR-post groups (Fig. 4B, Extended Data Fig. 7D). We next plotted the myeloid recovery signature (Fig. 2) in the UMAP space split by donor, pre-LVAD HF, U-post, and RR-post groups and observed a strong enrichment in the RR-post group and modest expression in the U-post group. The recovery signature was evident within all myeloid cell clusters (Fig. 4D). To functionally classify the recovery signature, we performed pathway analysis (EnrichR, WikiPathways database) and detected enrichment for EGF, HGF, androgen receptor, oncostatin M, and glucocorticoid receptor signaling (Extended Data Fig. 7C).

**Figure 4.**
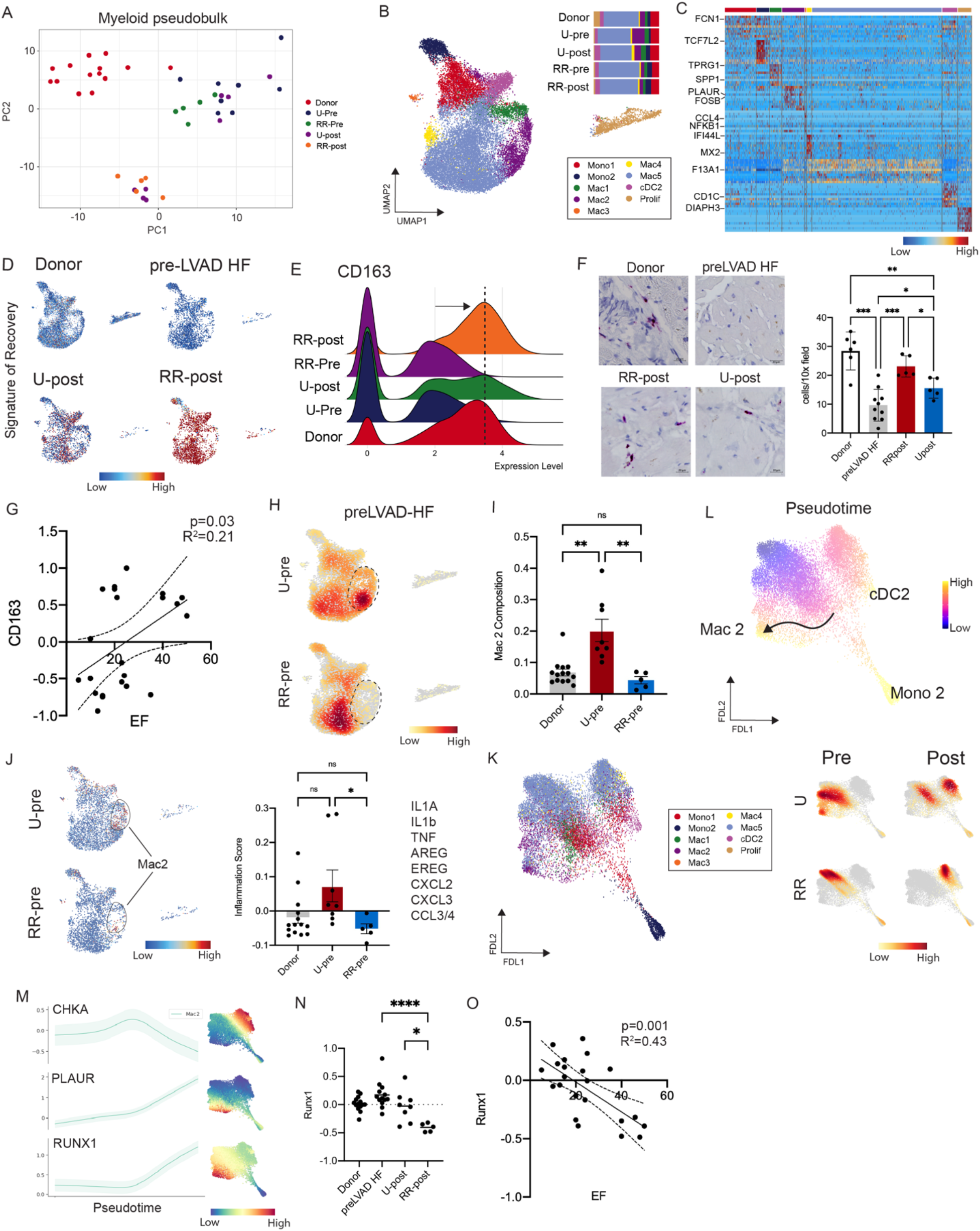
Pro-inflammatory macrophages and *RUNX1* diminished in reverse remodeling while resident macrophages show signs of recovery. (A) Myeloid pseudobulk PCA representation of each patient sample colored by condition. (B) UMAP embedding plot of myeloid cell states. (C) Heatmap of marker genes for distinct myeloid cell states. (D) Recovery up regulated signature in four conditions. (E) Ridge plot of CD163 expression split by five conditions. (F) RNAscope in situ hybridization of representative 10x fields from a donor, pre-LVAD HF, U-post, and RR-post sample (left) and quantification of number of CD163+ cells per 10x field across the four conditions (right). Brown-Forsythe and Welch ANOVA tests with multiple comparisons. Asterisks indicate degree of statistical significance. n = 26 total samples and Brown-Forsythe ANOVA test F = 20.78, DFn = 3, and p < 0.0001; donor vs preLVAD HF (***P = 0.0002), donor vs RR-post (nsP = 0.1251), donor vs U-post (**P = 0.0034), preLVAD HF vs RR-post (***P = 0.0001), preLVAD HF vs U-post (*P = 0.0256), and RR-post vs U-post (*P = 0.0108). (G) Linear regression of CD163 pseudobulk expression and patient ejection fraction in pre- and post-cohorts. Dotted line indicates 95% confidence interval, R^2^ goodness of fit, and p-value indicating whether the slope is significantly non-zero using an F test. (H) Gaussian kernel density estimation of cells in unloaded pre-LVAD and reverse remodeled pre-LVAD. (I) Mac2 cell state composition in donors and pre-LVAD patients. Dunn’s multiple comparisons test: donor vs U-pre (**P = 0.0045), U-pre vs RR-pre (**P = 0.0041), and donor vs RR-pre (nsP > 0.9999). (J) Gene set z-score for inflammation in unloaded pre-LVAD and reverse remodeled pre-LVAD (left) and quantified at patient level (right). Un-paired t-test with Welch’s correction: U-pre vs RR-pre (*P = 0.0321), donor vs U-pre (nsP = 0.1006), and donor vs RR-pre (nsP = 0.1602). (K) FDL embedding plot of myeloid cell states with colors from (B). (L) Palantir derived pseudotime in a FDL embedding plot with terminal states (Mac1, cDC2, and Mono2) using classical monocytes as a start state (top) and Gaussian density plot of cell number in FDL embedding split by four conditions (bottom). (M) CHKA, PLAUR, and RUNX1 gene expression along pseudotime (left) and expression in FDL embedding space (right). (N) Pseudobulk RUNX1 expression in pre-LVAD HF, U-post, and RR-post. Un-paired t-test with Welch’s correction: pre-LVAD HF vs U-post (nsP = 0.1177), pre-LVAD HF vs RR-post (****P < 0.0001), and U-post vs RR-post (**P = 0.0058). (O) Linear regression of RUNX1 pseudobulk expression and patient ejection fraction in pre- and post-cohorts. Dotted line indicates 95% confidence interval, R^2^ goodness of fit, and p-value indicating whether the slope is significantly non-zero using an F test.

We have previously identified a role for cardiac resident macrophages in adaptive remodeling of the heart^21–23^. To assess whether cardiac resident macrophages are involved in cardiac recovery, we generated RidgePlots and found that *CD163* (specific marker of cardiac resident macrophages) expression was decreased pre-LVAD HF groups compared to donor controls. Interestingly, *CD163* expression was restored to normal levels in the RR-post group while the U-post group displayed an intermediate phenotype (Fig. 4E). We validated our sequencing findings by performing *in situ hybridization* for *CD163* across patient groups. We observed a marked reduction in the number of CD163^+^ cells in pre-LVAD HF samples compared to donors. CD163^+^ cells increased to near normal levels in the RR-post group and modestly increased in the U-post group (Fig. 4F). To delineate whether *CD163* expression identifies a continuum of recovery, we performed a linear regression analysis of LV ejection fraction versus pseudobulk *CD163* expression at the patient level and found a modest correlation (R^2^=0.21, p=0.03) (Fig. 4G).

To dissect whether certain transcription cell states of myeloid cells predict recovery prior to LVAD implantation, we constructed a Gaussian kernel density plot of cell number in the pre-LVAD HF group split by U-pre and RR-pre conditions (Fig. 4H). Intriguingly, we found that the Mac2 cluster was overrepresented in the U-pre condition. Quantification of Mac2 composition across individual patients confirmed expansion of this population in the U-pre relative to donor and RR-pre groups (Fig. 4I, Extended Data Fig. 7D). Mac2 represented an inflammatory population that expressed *PLAUR* and several chemokines and cytokines (Fig. 4B, Extended Data Fig. 7A-B). To assess differences in inflammatory gene expression between U-pre and RR-pre groups, we constructed and plotted an inflammation gene set score (*IL1A, IL1B, TNF, AREG, EREG, CXCL2, CXCL3, CCL3, CCL4*) in UMAP space. The inflammatory signature was localized to the Mac2 population and present selectively in the U-pre group (Fig. 4J), suggesting that the presence of inflammatory macrophages is a negative predictor of cardiac recovery.

To decipher the origin of Mac2, we performed pseudotime analysis using Palantir in the HF samples and identified three terminal monocyte-derived states: Mac2, cDC2, and Mono2 (Fig. 4K-L). Cell density plots showed strong phenotype shifts between each pre-LVAD and post-LVAD group when viewed in a force directed layout (FDL) space (Fig. 4L). Consistent with the above findings, we observed an enrichment for Mac2 in the U-pre group. Notably, post-LVAD implantation RR-post converged towards a Mac5 phenotype while U-post displayed a divergence towards Mac5 and Mac2, 4 (Fig. 4B,L). In particular, the RR-post group converged towards a phenotype marked by enriched *CHKA* expression (Fig. 4M). This population largely consistented of *CD163^+^* cardiac resident macrophages. We next plotted MAGIC imputed gene expression for *CHKA, PLAUR,* and *RUNX1* along pseudotime within the Mac2 lineage. We observed a monotonic increase in *PLAUR* and *RUNX1* expression and decreased *CHKA* expression as cell differentiated towards Mac2 (Fig. 4M), highlighting competing differentiation trajectories between heart failure and recovery.

To dissect regulatory changes that may underlie transcriptional signatures of heart failure and recovery in myeloid cells, we performed TF enrichment analysis with DoRothEA^24^ in the U-post and RR-post groups. Transcription factors associated with inflammation (*RUNX1, NFKB1, NFKB2, STAT3*, *ATF2, JUN*) were enriched in myeloid cells from the U-post group (Extended Data Fig. 7E). Quantification of *RUNX1* expression at the patient level via pseudobulk analyses revealed increased *RUNX1* expression in the donor, pre-LVAD HF, U-post groups. The expression of *RUNX1* was markedly diminished in the RR-post group relative to all other groups (Fig. 4N) – these results suggest that *RUNX1* downregulation may modulate macrophage phenotype towards a unique state unlike a healthy or failing heart. We next performed a linear regression for LV ejection fraction versus *RUNX1* pseudobulk expression and found a strong negative correlation (R^2^=0.43 and p=0.001) (Fig. 4O). To assess the interplay between *RUNX1* and recovery we plotted a pseudobulk heatmap of *RUNX1* target genes that were differentially expressed between U-pre and RR-pre groups and identified numerous pro-inflammatory mediators that were enriched in the U-pre group. Collectively, these analyses highlight the possibility that *RUNX1* may prevent recovery by promoting pro-inflammatory gene expression in macrophages (Extended Data Fig. 7F).

### Cardiac fibroblast remodeling in cardiac recovery

Cardiac recovery is associated with reductions in myocardial fibrosis^25, 26^. To ascertain how cardiac fibroblasts shift during recovery, we first performed a pseudobulk PCA. Donor samples clustered separately from the pre-LVAD and post-LVAD groups. A modest separation was also observed between the RR-post group and the pre-LVAD HF groups (Fig. 5A). We then sub-clustered the fibroblasts into 8 cell states (Fig. 5B) marked by distinct transcriptional signatures (Fig. 5C, Extended Fig. 8A, B). Cell composition and density analysis demonstrated that Fib1 (*SCN7A*) was enriched in donors, while Fib3 (*POSTN, THBS4, MEOX1*) and Fib7 were enriched in HF (*GPC6*) (Fig. 5D). Pathway analysis revealed enrichment in genes associated with extracellular matrix remodeling in Fib3 and actin binding in Fib7 (Extended Data Fig. 8C). To assess whether recovered fibroblasts represent a reversion to the normal state, we generated a heat map of genes enriched in donor control and HF fibroblasts at the pseudobulk level. This analysis revealed the presence of both persistently dysregulated (*SVEP1, FAP, POSTN, GPX3, APOD*) and normalized genes (*MEOX1, TGFBR3, ACSM3, PID1*) in recovered fibroblasts (Fig. 5E). Consistent with these findings, *in situ* hybridization for *POSTN* revealed persistence of this fibroblast population during recovery (Fig. 5O). We then generated RidgePlots of the fibroblast specific recovery signature and detected an enrichment within the RR-post group (Fig. 5F-H). Gene ontology of the top genes upregulated in recovery suggested associations with cytoskeletal organization, glucose homeostasis, and receptor tyrosine kinase signaling (Fig. 5G). Conversely, the top pathways downregulated in recovery included TNF-a/NF-kB and TGF-beta signaling (Fig. 5I).

**Figure 5.**
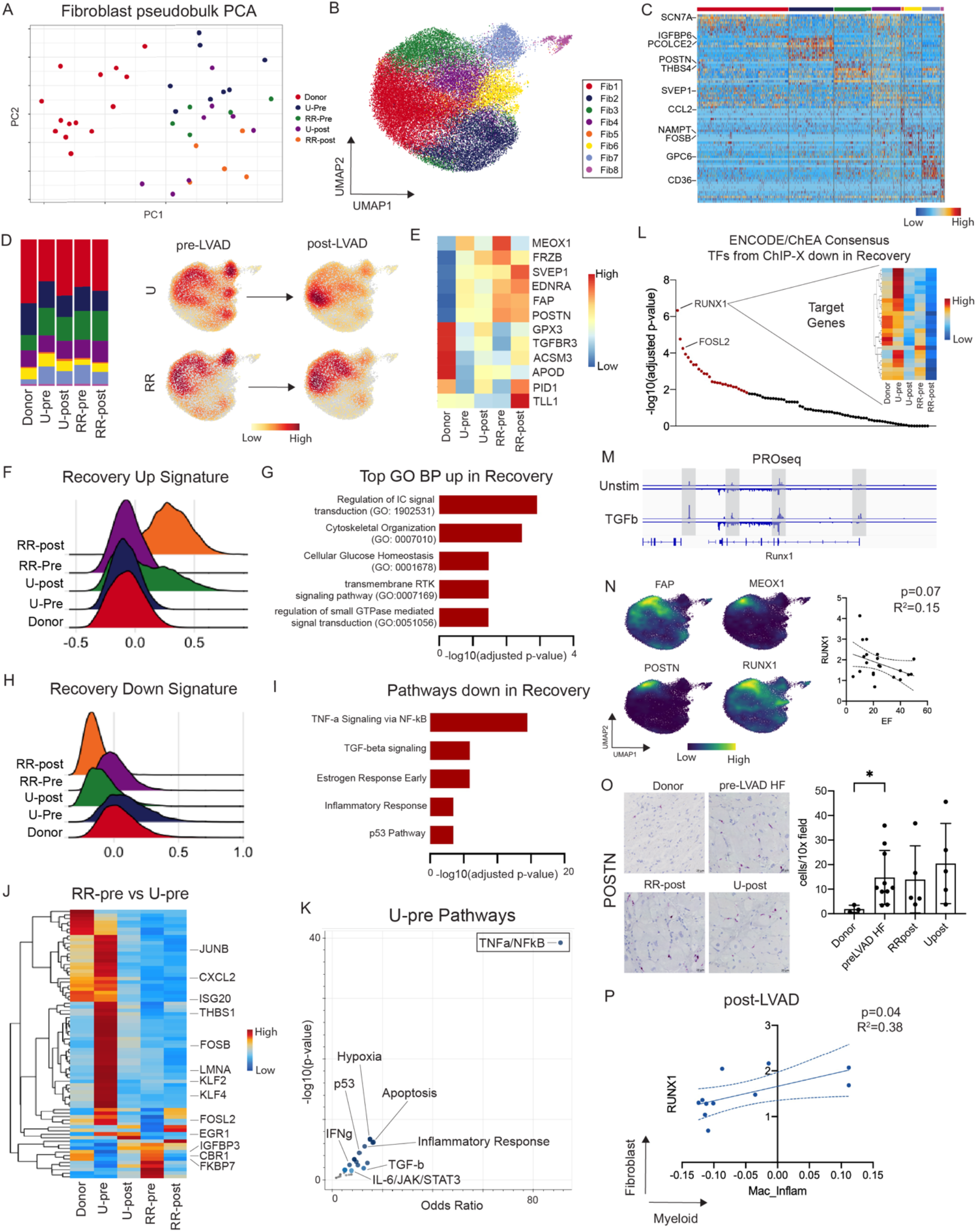
RUNX1 is downregulated in fibroblasts in cardiac recovery. (A) Fibroblast pseudobulk principle component analysis representation of each patient sample colored by condition. (B) UMAP embedding plot of fibroblast cell states. (C) Heatmap of marker genes for distinct fibroblast cell states. (D) Cell composition of fibroblast cell states across five conditions (left) and Gaussian kernel density estimation of cells in four conditions pre- and post-LVAD implantation (right). (E) Heatmap of expression of canonical genes up and downregulated in fibroblasts in heart failure across five conditions. (F) Up and (H) down regulated pseudobulk recovery signature Ridge plot split across five conditions. (G) GO biological processes pathways enriched in recovery and (I) pathways down in recovery. (J) Heatmap of pseudobulk DE analysis between RR-pre and U-pre split across 5 conditions. (K) Volcano Plot of pathways enriched in unloaded group pre-LVAD implantation where each point is a gene set pathway, and the color of the points represents degree of statistical significance. (L) Transcription factors down in cardiac recovery from ENCODE/ChEA consensus; x-axis is transcription factors and red dots are TFs which are statistically significant. Heatmap showing expression of RUNX1 target genes across five conditions. (M) PRO-seq coverage in unstimulated and TGF-b treated *in vitro* fibroblasts (GSE15582) at the *RUNX1* locus. (N) Density plots of HF genes in UMAP embedding (left) and linear regression of RUNX1 pseudobulk expression and patient ejection fraction in pre- and post-cohorts (right). Dotted line indicates 95% confidence interval, R^2^ goodness of fit, and p-value indicating whether the slope is significantly non-zero using an F test. (O) RNAscope in situ hybridization of representative 10x fields from a donor, pre-LVAD HF, U-post, and RR-post sample (left) and quantification of number of POSTN+ cells per 10x field across the four conditions (right). Brown-Forsythe and Welch ANOVA tests with multiple comparisons. Asterisks indicate degree of statistical significance. Donor vs pre-LVAD HF (*P = 0.0272). (P) Linear regression of RUNX1 pseudobulk expression in fibroblasts and inflammatory signature in macrophages in post-LVAD cohort. Dotted line indicates 95% confidence interval, R^2^ goodness of fit, and p-value indicating whether the slope is significantly non-zero using an F test.

To identify cardiac fibroblast transcriptional changes that predict recovery, we performed pseudobulk differential expression analysis between the RR-pre and U-pre groups. We detected multiple robust differences between groups including multiple genes association with inflammation (*JUNB, CXCL2, KLF2, KLF4, ISG20, FOSL2*) increased in the U-pre group (Fig. 5J). Pathway analysis of dysregulated genes showed strong upregulation of TNFa/NF-kB, inflammatory response, TGFb, IFNg, and IL-6/STAT3 pathways (Fig. 5K).

To explore transcriptional programs that may drive recovery, we examined ENCODE/ChEA Consensus transcription factors from the ChIP-X database and identified *RUNX1* as the most downregulated transcription factor during recovery. We then plotted a heatmap of genes predicted to be regulated by *RUNX1* during recovery and identified a strong downregulation in the RR-post group (Fig. 5L). Given the above changes observed in *RUNX1* expression, predicted activity, and TGF-b enrichment, we leveraged published PRO-seq data from *in vitro* fibroblasts stimulated with TGF-b to assess the level of active transcription around the *RUNX1* locus. We found increased RNA polymerase activity at the *RUNX1* locus following TGF-b stimulation relative to a unstimulated control (Fig. 5M), suggesting that TGF-b signaling may regulate *RUNX1* expression in cardiac fibroblasts. Next, we sought to delineate *RUNX1* expression across fibroblast cell states. Density plots revealed that *RUNX1* was expressed in Fib3 along with genes known to contribute to myocardial fibrosis including *POSTN, FAP,* and *MEOX1*^18, 27, 28^. Linear regression of ejection fraction versus pseudobulk *RUNX1* expression showed a negative correlation (R^2^=0.15 and p=0.07) (Fig. 5N). To explore a link between macrophage inflammatory gene expression and associated *RUNX1* expression in fibroblasts, we performed a linear regression of fibroblast specific *RUNX1* expression vs the macrophage inflammatory score (Fig. 4N) and observed a correlation at the patient level (R^2^=0.38 and p=0.04) (Fig. 5P). These results suggest that elevated inflammation in macrophages in U-pre relative to RR-pre may hinder cardiac recovery by modulating *RUNX1* expression in fibroblasts.

### RUNX1 gene regulatory network is predictive of recovery

To decipher the potential contribution of *RUNX1* towards promoting the recovery phenotype, we performed the following analyses. First, we validated changes in *RUNX1* expression in stromal cells across groups using *in situ* hybridization. Quantitation of the number of *RUNX1^+^*stromal cells per 10x field, revealed that there were greater numbers of *RUNX1^+^*cells in the pre-LVAD HF and U-post groups relative to donors. The RR-post group displayed reduced numbers of *RUNX1^+^* cells relative to the pre-LVAD HF groups and was comparable to donors (Fig. 6A-B). Next, we assessed whether *RUNX1* target gene expression is predictive of cardiac recovery using deep learning. We split the pre-LVAD HF group (U-pre and RR-pre) into a training and test set, built and trained a Keras classifier model, and applied the model in the test set to predict whether cells from recovered samples were derived from the U-pre or RR-pre patients (Fig. 6C). We also trained a random forest classifier as a comparator to our Keras model. We found that both the Keras model and random forest classifier predicted recovery. The Keras model had a higher AUC than the random forest classifier in predicting recovery (myeloid: Keras=0.945, RF=0.789 and fibroblast: Keras=0.942, RF=0.836) (Fig. 6D-E). We then tested whether perturbation of *RUNX1* could facilitate recovery in human macrophages and fibroblasts. We utilized CellOracle^17, 29^ to build cell specific gene regulatory networks and performed an *in silico* deletion of *RUNX1*. To define directional changes in cell fate resulting from *RUNX1* perturbation, we constructed a vector field and visualized the data UMAP space (Fig. 4B, Fig. 5B). We leveraged Palantir pseudotime trajectory analysis to infer the baseline flow of cells between states (Extended Data Fig. 9A-B). To identify cell states enriched and depleted following *RUNX1* pertubation, we took the inner product between control and experimental vector fields. This analysis indicated that *RUNX1* perturbation in myeloid cells resulted in movement of cells away from the Mac1 and Mac2 states and towards the Mac5 and monocyte states, signifying a predicted block in the differentiation of inflammatory macrophages and preservation of cardiac resident macrophages (Fig. 6F-H). *RUNX1* perturbation in fibroblasts resulted in movement of cells away from the Fib3 and Fib6 states and towards the Fib0, Fib2, Fib7 states, suggesting a shift away from states associated with fibrosis and heart failure (Fig. 6G-I). These findings indicate that *RUNX1* expression is reduced during recovery and predicts that downregulation of *RUNX1* activity in macrophages and fibroblasts signifies the potential for recovery and drives shifts in macrophages and fibroblasts away from states associated with heart failure.

**Figure 6.**
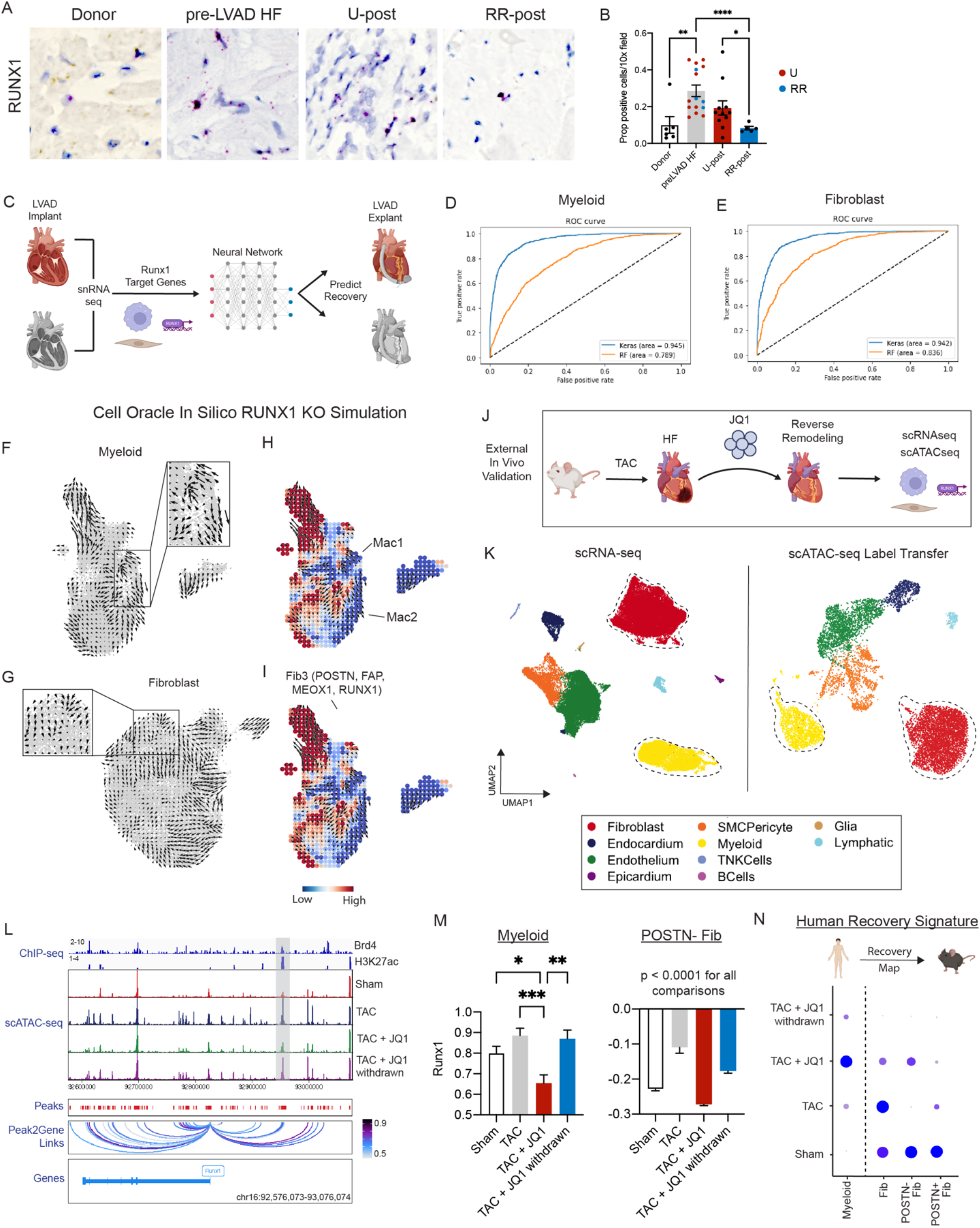
*RUNX1* perturbation *in silico* and *in vivo* facilitates cardiac recovery. (A) RNAscope in situ hybridization for Runx1 in a representative donor, pre-LVAD HF sample, U-post, and RR-post. (B) RNAscope images quantified as proportion of positive interstitial nuclei/total interstitial nuclei per 10x field. Brown-Forsythe and Welch ANOVA test comparing each condition to donor. Asterisks indicate degree of statistical significance. n = 37 total samples and Brown-Forsythe ANOVA test F = 7.686, DFn = 3, and p = 0.0009. Asterisks indicate degree of statistical significance: *P < 0.05, **P < 0.01, and ***P < 0.001. p-value specified for non-significant comparison. (C) Schematic of machine learning approach used to predict recovery in macrophages and fibroblasts with Runx1 target genes as features. ROC curves with accuracy metrics for test dataset in predicting recovery in (D) macrophages and (E) fibroblasts using a Keras deep neural net classifier model and random forest classifier. CellOracle in silico Runx1 KO simulation quiver plot of vector field in (F) macrophages and (G) fibroblasts. Perturbation score with vector field in (H) macrophages and (I) fibroblasts. (J) Study design of external validation dataset. (K) scRNA-seq and scATAC-seq UMAP embedding with cell labels from RNA-seq label transferred onto ATAC-seq dataset using publicly available data (GSE15582). (L) Mouse *Runx1* locus showing from top to bottom: ChIP-seq for BRD4 (GSE46668) and H3K27ac (ENCSR000CDF) in adult mouse heart and scATAC-seq in mouse fibroblasts split by four conditions with called peaks (GSE15582), and peak2gene links from ArchR. Numbers above tract indicate ranges of normalized tag densities. Highlighted area is a peak linked to the Runx1 gene. (M) *Runx1* expression in macrophages (left) and *Postn-*fibroblasts (right) in Sham, TAC, TAC + JQ1, and TAC + JQ1 withdrawn. Brown-Forsythe and Welch ANOVA multiple comparisons test. Myeloid: Sham vs TAC + JQ1 withdrawm (*P = 0.0492), TAC vs TAC + JQ1 (***P = 0.0003), and TAC + JQ1 vs TAC + JQ1 withdrawn (**P = 0.0021). Fibroblast: ****P < 0.0001 for all pairwise comparisons. (N) Dotplot of Human myeloid and fibroblast cardiac recovery gene signature plotted in mouse macrophages in fibroblasts split by 4 conditions.

To evaluate the biological plausibility of the above predictions, we leveraged published *in vivo* scRNA-seq and scATAC-seq data obtained from a study investigating the potential of BRD4 inhibition (JQ1 treatment) to promote recovery in a mouse model of pressure overload heart failure (Transverse Aortic Constriction, TAC)^18, 27^ (Fig. 6J). We re-processed the scRNA-seq data and annotated major cell populations, independently clustered the scATAC-seq data, and performed label transfer to annotate scATAC-seq clusters using the RNA information (Fig. 6K). We selected the fibroblasts from the ATACseq dataset to explore changes in chromatin accessibility around the *Runx1* locus with JQ1 treatment. We then performed peak to gene linkage around the *Runx1* locus and identified several regions with increased accessibility in TAC relative to sham fibroblasts that decreased with JQ1 treatment, and subsequently increased following withdrawal of JQ1. Using publicly available BRD4 and H3K27ac ChIP-seq data from mouse left ventricle we validated many of these peaks as active enhancer marks that were binding site for BRD4 (Fig. 6L). Next, we quantified *Runx1* expression from the scRNA-seq data in macrophages and fibroblasts and found that JQ1 treatment was associated with reduced *Runx1* expression (Fig. 6M). Finally, to assess the relationship between JQ1 treatment and human signatures of recovery, we examined the expression of human macrophage and fibroblast genes associated with recovery across the mouse conditions. Within macrophages, the human recovery signature was enriched in the JQ1 treated group and not the sham group indicating that BRD4 inhibition drives the acquisition of the recovered state. The transcriptional signature of recovery was evident in both sham and JQ1 treated conditions highlighting that BRD4 inhibition promotes both the acquisition of the recovered state and reversion to baseline in cardiac fibroblasts (Fig. 6N).

## Discussion

By performing snRNA-seq on donor hearts and samples from heart failure patients who either recovered or did not recover LV systolic function following LVAD implantation, we defined the single cell landscape of human cardiac recovery. This work elucidated that the recovered heart represents a distinct biological entity rather than simply a reversion to the normal or non-failing state. We discovered that the transcriptional signature of cardiac recovery is uniquely encoded across cell types with dominant contributions from cardiac macrophages and fibroblasts. Pro-inflammatory macrophage and fibroblast states were found to be negative predictors of recovery. Construction of gene regulatory networks predicted that downregulation of RUNX1 transcriptional activity in cardiac macrophages and fibroblasts as a central mechanism orchestrating cardiac recovery, a possibility supported by machine learning and *in silico* transcriptional factor perturbation techniques and corroborated in a mouse model of cardiac recovery mediated by BRD4 inhibition These findings highlight the possibility that inhibition of RUNX1 may facilitate cardiac recovery.

For patients suffering from heart failure, cardiac recovery represents the most ideal outcome. Recovery of LV systolic function is observed in select individuals following the initiation of anti-remodeling medications (beta adrenergic, angiotensin II and neprilysin inhibitors, aldosterone antagonists) and mechanical unloading. The frequency of cardiac recovery declines with the severity of heart failure. Individuals with end-stage heart failure that are eligible for LVAD implantation and heart transplantation display the lowest incidence of recovery (1-2%)^3–6^. It has been postulated that combined mechanical unloading and pharmacologic treatment may serve as an approach to increase recovery rates in the growing advanced heart failure population where treatment strategies remain limited by recipient eligibility for heart transplantation and donor organ availability^30, 31,32,33^. An improved understanding of the mechanistic basis by which hemodynamic changes imposed by mechanical unloading translate to molecular and structural changes associated with cardiac recovery^34, 35^ is essential for the development of new therapeutic strategies to increase the frequency of cardiac recovery. Thus, it is of the utmost clinical importance to delineate the cell types and mechanisms that orchestrate acquisition and maintenance of cardiac recovery.

The concept that recovery is not merely a reversion to normal is supported by both clinical and biological data. Clinical trials investigating outcomes following withdrawal of anti-remodeling therapy and mechanical unloading in heart failure patients who recovered LV systolic function identified a significant rate of recurrent heart failure and deterioration of cardiac function. These data suggest that cardiac recovery represents a state of remission and indicate that maintenance of cardiac recovery requires ongoing intervention^36, 37^. The possibility that the recovered heart is not normal is also supported by physiological and pathological investigation demonstrating abnormal contractile reserve, blunted force frequency relationship, and persistent fibrosis^4, 9, 38–42^. Consistent with our findings, bulk RNA sequencing in a mouse model of cardiac recovery triggered by mechanical unloading identified a transcriptional signature that was distinct from both baseline and failing conditions. While some heart failure genes normalized, the majority remained persistently dysregulated and new signatures associated with recovery emerged^43^. By generating a cell type specific transcriptional map of human cardiac recovery, we similarly observed that the recovered human heart retains signatures of heart failure, and instead, found that recovery was associated with the emergence of cell specific transcriptional states that were not found in healthy or diseased conditions.

Intriguingly, we found that transcriptional signatures of cardiac recovery were predominately encoded within stromal cells with the largest contribution from cardiac macrophages. Within this compartment, we detected distinct contributions from monocyte-derived and cardiac resident macrophages, populations with opposing functions and effects on heart failure pathogenesis^21, 44, 45^. Patients who did not recover displayed an enrichment in monocyte-derived macrophages expressing genes implicated in myocardial inflammation and heart failure (*PLAUR, IL1B, TNF, CCL4)*^13^. The presence of this population was a negative predictor of recovery. These findings are consistent with prior work identifying an association between the abundance pro-inflammatory monocyte-derived macrophages that expressed CCR2 and failure to recover^44^. In contrast, we observed that CD163^+^ cardiac resident macrophages are lost in heart failure and revert to normal levels in those who recover. Given that cardiac resident macrophages strongly express transcriptional signatures associated with cardiac recovery and are known to possess functions important for tissue healing and remodeling^23, 46, 47^, it is likely that this population is a central mediator of the recovery process.

Cardiac fibroblast activation is an important mechanism of myocardial fibrosis and heart failure pathogenesis^28, 48^. The exact role of fibroblasts in cardiac recovery continues to be debated. Within the recovered heart, cardiac fibroblasts retained signatures of fibroblast activation including the expression of *FAP, MEOX1,* and *POSTN*^13, 14, 18, 28^. Interestingly, cardiac recovery was associated with reduced expression of genes associated with inflammatory and TGF-b signaling in cardiac fibroblasts, markers of immune cell driven fibroblast activation^18, 49^. When viewed together with the observation that pro-inflammatory macrophages persist in patients that do not recover, these findings suggest that resolution of inflammatory signaling between macrophages and fibroblasts may be essential for recovery. These findings are consistent with the paradigm that para-inflammation leads to persistent tissue dysfunction whereas resolution of inflammation restores ssue function^50^.

Prior studies have expanded on the role of the cardiac microenvironment and cellular crosstalk in modulating cardiac recovery^51^. To unravel transcriptional networks along the macrophage-fibroblast axis driving recovery, we performed transcription factor enrichment and constructed gene regulatory networks associated with recovery. This analysis identified downregulation of RUNX1 activity as a key feature of recovery. *RUNX1* is a transcription factor with important roles in inflammatory responses and hematological cancers^52, 53^. Studies performed in zebrafish suggest a role in fibroblast activation^54, 55^. To explore a relationship between *RUNX1* in macrophages and fibroblasts in cardiac recovery, we applied a deep learning approach^56, 57^ and found that RUNX1 target gene expression measured at the time of LVAD implantation predicts acquisition of recovered versus non-recovered states in both cell populations. We then used CellOracle^17^ to build macrophage and fibroblast specific gene regulatory networks and ascertained the predicted effects of *RUNX1* deletion using *in silico* transcription factor perturbation. Within macrophages, *RUNX1* perturbation resulted in predicted loss of pro-inflammatory macrophages and pathogenic fibroblasts with shifts towards states observed in recovered hearts. We then tested our predictions in a clinically relevant mouse model of cardiac recovery mediated by JQ1 treatment, an inhibitor of BRD4^18^. Using publicly available multi-omic data, we showed that BRD4 binds to the *Runx1* enhancer in mouse hearts^27^, suggesting that inhibition of BRD4 may disturb the *Runx1* gene regulatory network. We identify enhancer peaks in fibroblasts linked to the *Runx1* gene that display increased accessibility in heart failure that are diminished by JQ1 treatment, similarly to what was observed at the *Meox1* locus^18^. Consistent with disruption of RUNX1 activity, JQ1 treatment led to reduced *Runx1* expression in macrophages and fibroblasts. Finally, we show that JQ1 treatment was sufficient to trigger the emergence of the human cardiac recovery signature in macrophages and fibroblasts. Collectively, these findings highlight the possibility that RUNX1 inhibition may serve as an approach to facilitate cardiac recovery.

Our study is not without limitations. While our sample size is limited, availability of paired (pre- and post-LVAD implant) myocardial tissue specimens from patients who recover is incredibly scarce given the low incidence of cardiac recovery and small numbers of clinical centers with robust pipelines to phenotype and capture this unique patient population. It is important to note that the *in silico* approach used to perturb *Runx1* within macrophages and fibroblasts does not consider cellular crosstalk. While an important limitation, CellOracle transcription factor perturbation studies have been validated *in vitro* and *in vivo* and used by other groups^17, 29, 58^. We recognize that there is no perfect mouse model to recapitulate cardiac recovery^43^ and the pressure overload JQ1 mediated recovery dataset has inherent limitations. This dataset is used to test human derived hypothesis in conjunction with the human recovery data to offset this issue. Lastly, while we show *in silico* perturbation of *Runx1* is predicted to facilitate the recovery phenotype, we recognize that conditional deletion of *Runx1* in appropriate models is necessary to establish causality.

In conclusion, we provide a comprehensive single cell transcriptomic map of human cardiac recovery, establish that cardiac recovery is a biological state distinct from healthy and disease, and unravel cell type specific signatures of recovery. Furthermore, we identify shifts in macrophage and fibroblast phenotypes driven by a *RUNX1* gene regulatory network that predict the propensity for recovery. Finally, we provide orthogonal sources of evidence to suggest that disruption of *RUNX1* in macrophages and fibroblasts drives the recovered phenotype and that RUNX1 inhibition may be an effective approach to facilitate cardiac recovery.

## Acknowledgments

KL is supported by the Washington University in St. Louis Rheumatic Diseases Research Resource-Based Center grant (NIH P30AR073752), the National Institutes of Health [R01 HL138466, R01 HL139714, R01 HL151078, R01 HL161185, R35 HL161185], Leducq Foundation Network (#20CVD02), Burroughs Welcome Fund (1014782), and Children’s Discovery Institute of Washington University and St. Louis Children’s Hospital (CH-II-2015-462, CH-II-2017-628, PM-LI-2019-829), Foundation of Barnes-Jewish Hospital (8038-88), and generous gifts from Washington University School of Medicine. JMA is supported by the American Heart Association Predoctoral Fellowship (826325). PM is supported by American Heart Association Postdoc Fellowship (916955). Figure 1A and 6C were created in BioRender.com. We thank the Genome Technology Access Center at the McDonnell Genome Institute at Washington University School of Medicine for help with genomic analysis. The Center is partially supported by NCI Cancer Center Support Grant #P30 CA91842 to the Siteman Cancer Center. This publication is solely the responsibility of the authors and does not necessarily represent the official view of NCRR or NIH.

## Author Contributions

SD contributed to LVAD sample acquisition and clinical phenotyping. PM, LL, AB, and AK isolated nuclei for snRNA-seq. JA performed computational analysis. JA, KK, and SM performed GRN analysis and in silico TF perturbation analysis. JA, LL, PM, LS, DS, FK, and BK performed RNA in situ hybridization and immunohistochemistry experiments and analyzed and processed images. JA, FL, RK, DM, SD, and KL assisted in the interpretation of the data. JA constructed all figures and JA and KL drafted the manuscript. KL is responsible for all aspects of this manuscript including experimental design, data analysis, and manuscript production. All authors approved the final version of the manuscript.

## Materials and Methods

### Ethical Approval for Human Specimens

This study complies with all ethical regulations associated with human tissue research. Acquisition of donor samples was approved by the Washington University Institutional Review Board (study no. 201104172). All samples were procured with informed consent from patients obtained by Washington University in St Louis School of Medicine. No compensation was provided for participation. Donor and patient demographical details can be found in Supplemental Table 1. Cardiac phenotyping was performed at the time of LVAD implantation and explant and relevant cardiac function metrics are provided in Supplemental Table 2. Clinical baseline data was collected for all patients at the time of LVAD implantation and is provided in Supplemental Table 3.

### Sample Selection clinical phenotyping

The recovery cohort of patients were selected to match for the following variables to the best extent: EF at the time of LVAD implant, sex, age, and clinical risk profile. Age and sex matched donors were then pulled from the Washington University in St Louis School of Medicine biobank repository. Within the LVAD cohort, patients were assigned as “reverse remodeled” or unloaded”. Donors were selected to age and sex match with the LVAD samples. RR and U samples were chosen such that EF at the time of LVAD implantation is not different between the two groups.

### Human single nuclei isolation and library preparation snRNA-seq

Cardiac tissues from LVAD cores at the time of LVAD implant (U/RR-pre) and adjacent to core samples at the time of explant (U/RR-post) from paired patient were flash frozen using liquid nitrogen. Identical regions from the apex of LV from explanted donors were used. Single nuclei suspensions were generated as previously described. In brief: flash frozen sections were minced with a razor blade, transferred to a Dounce Homogenizer containing 1 mL of lysis buffer (10 mM Tris-HCl, pH 7.4, 10 mM NaCl, 3 mM MgCl_2_ and 0.1% NP-40 in nuclease-free water) on ice. Samples were homogenized using five strokes, an additional 1 mL of lysis buffer added, and incubated on ice for 15 mins. Samples were then filtered with a 40μm filter and filter was rinsed with 1mL of lysis buffer. The mixture was then centrifuged at 500*g* for 5 min 4 °C, resuspended in 1mL nuclei wash buffer (2% BSA and 0.2 U μl^−1^ RNase inhibitor (Thermo Fisher, cat. no. AM2694) in 1× PBS) and, filtered using a 20*μ*m pluristrainer (Pluriselect, cat. No. SKU43-50020-03). Filtered solution as centrifuged using the above criteria and resuspended in 300 μL Nuclei Wash Buffer and transferred into a 5mL tube for flow cytometry. Subsequently, 1 μl DRAQ5 (5 mM solution; Thermo Fisher, cat. no. 62251) was added, sample gently vortexed, and allowed to incubate for 5 min prior to sorting. DRAQ5^+^ nuclei were sorted into 300 μL Nuclei Wash Buffer using a BD FACS Melody (BD Biosciences) with a 100 µM nozzle. Sorted nuclei were then centrifuged using the above conditions and resuspended in Nuclei Wash Buffer for a final target concentration of 1,000 nuclei/μL – nuclei were counted on a hemocytometer. Based on the nuclei concentration, 10,000 target nuclei were loaded onto a Chip G for GEM generation using the Chromium Single Cell 5ʹ Reagent v1.1 kit from 10X Genomics. Reverse transcription, barcoding, complementary DNA amplification and purification for library preparation were performed as per the Chromium 5ʹ v1.1 protocol at the McDonnel Genome Institute. Sequencing was performed on a NovaSeq 6000 platform (Illumina) at a target read depth of 100,000 at the McDonnel Genome Institute.

### Global transcriptomic map generation

Nuclei fastq files were aligned to the whole genome pre-mRNA reference generated from the GRCh38 transcriptome (with the intron flag included) using CellRanger v3 (10X Genomics). Nuclei were filtered to include those with 1000 < RNA UMI count < 10,000 and mitochondrial reads < 5%. After initial QC, scrublet was ran on each sample separately in Python with default parameters to score nuclei and nuclei with a doublet score > 0.2 were excluded from downstream analysis. We then leveraged supervised doublet removal as previously described on a per cell type basis before combining all objects. Briefly, clusters were annotated into major cell populations, each major cell type was subsetted, and re-normalized, PCA, UMAP embedding, clustering, and DE analysis performed. Sub-clusters which did not express the gene signature of the cell type or had overlapping genes across different cell types were removed. Post-contamination, all cell type objects were merged to construct a cleaned object. After QC and doublet removal downstream analysis was performed in Seurat v4^59^. The cleaned object was normalized using SCTransform^60^ with regressing out mitochondrial percent and RNA UMI counts. We then computed the principal components and used these to integrate all samples with harmony^61^. Informed by the ElbowPlot, we used 80 components to construct the UMAP embedding, find nearest neighbors, and clustered the data at multiple resolutions. We then used the FindAllMarkers function in Seurat to perform differential expression testing and annotated clusters into distinct cell types based on canonical gene markers. To define cell states within each major cell type, we subsetted the major cell populations, re-normalized, re-clustered, re-computed the UMAP, and annotated cell states. Both the global object and each cell type object was saved and used for downstream analysis and plotting in Seurat. All differential gene expression to identify cell types or cell states was performed using the normalized assay with the FindAllMarkers function and the Wilcoxon rank sum test with a min.pct = 0.1 and logfc.threshold = 0.25. A heatmap of the top10 genes was made across cell types/states. To further substantiate our annotations, we used the top genes in each cell type/state to create a gene set z-score to see separation across clusters.

### Pseudobulk Differential Gene Expression

Pseudobulk differential gene expression was performed using the DESeq2^62^ package. After QC, cells were subsetted for each cell type, raw counts extracted, raw counts were aggregated to the sample level, data normalized using a regularized log transform, a pseudobulk PCA performed, and differential expression analysis between conditions of interested via DESeq2. For pseudobulk DE analysis, we made the following comparisons: (1) pre-LVAD HF (U-pre and RR-pre) vs donor, (2) RR-post vs donor, and (3) RR-post vs pre-LVAD HF (U-pre and RR-pre). Genes were deemed statistically significant if adjusted p-value < 0.05 and absolute(log2FC) > 0.58. Statistically significant DE genes from comparisons (1) – (3) were then used to compute cardiac recovery, persistent HF, and HF reversed genes within R as defined by the Venn Diagrams. Additionally, we constructed UpSet plots to look at gene overlap across different cell type permutations.

### Continuum analysis with ejection fraction

To connect transcriptional changes with patient ejection fraction within the LVAD cohort we performed a simple linear regression of EF versus: (1) pseudobulk expression of genes of interest in specific cell types such as CD163 and RUNX1 and (2) cell type specific pseudobulk gene set score of recovery up-and down-regulated genes. Pseudobulk aggregation was done at the sample level as defined above. Regressions were performed in Prism 9 (Graphpad), an R^2^ computed for goodness of fit, and a p-value indicating whether the slope is non-zero (F-test).

### Cell composition and density shift calculations

We used R to compute cell type composition across conditions. To assess shifts in cell density within both the global object and within individual cell types, we converted the .rds object to a .h5ad file format and used scanpy.tl.embedding function which employs a Gaussian kernel density estimation of cell number within the UMAP embedding. Density values are scaled from 0-1 within that category.

### RNAscope In Situ Hybridization

Flash frozen LV samples were fixed for 24 hr at 4 °C in 10% neutral buffered formalin, washed in 1× PBS, and embedded in paraffin. Paraffin-embedded sections were cut at an 8 μm thickness using a microtome. RNA In situ hybridization was performed using the RNAScope^63^ Multiplex Fluorescent Reagent kit v2 Assay, RNAScope 2.5 HD Detection Reagent as per the protocol – RED and RNAScope 2.5 HD Duplex Assay kits (Advanced Cell Diagnostics, ACDBio) using probes designed by Advanced Cell Diagnostics. Fluorescent images were collected using a Zeiss LSM 700 laser scanning confocal microscope. The following RNAScope probes from ACDBio were used: MYH6 (555381), NPPA (531281), CD163 (417061), POSTN (409181), PLAUR, and RUNX1. Chromogenic/brightfield/fluorescent7 images were acquired using a Zeiss Axioscan Z1 automated slide scanner. Image processing was performed using Zen Blue and Zen Black (Zeiss). Positive cells were counted using either of two approaches: (1) For fluorescent images the number of positive cells counted per 10x field or (2) for chromogenic images number of positive interstitial cells/total number of interstitial nuclei per 10x field. For CD163 and POSTN, donor and DCM samples were used from our previously published study and re-quantified as per the above approach to ensure comparability.

### Pseudotime analysis

Palantir^64^ was used to perform pseudotime analysis. To dissect monocyte fate specification in HF and cardiac recovery we subsetted the myeloid object to include U-pre, U-post, RR-pre, and RR-post and excluded the proliferating cells. The normalized count matrix was then exported as a .txt file and loaded into Python. Palantir was then used to compute a PCA, diffusion map, and a multiscale low dimensional embedding computer with 5 eigenvectors. A force directed layout was then computed for visualization of trajectories and MAGIC was used to impute data for visualization. The Palantir simulation was executed with classical monocytes as the starting state, no terminal states specified, and 600 waypoints used. Generalized Additive Models within Palantir was then sued to compute gene expression trends along the Mac2 lineage using the plot_gene_trend_heatmaps function. Pseudotime, entropy, and FDL embedding values were exported and subsequent plotting for visualization was performed in R/Seurat.

### Pathway Analysis and transcription factor enrichment

Statistically significant DE cardiac recovery genes as defined above from the pseudobulk section were used to perform pathway enrichment analysis. EnrichR (https://maayanlab.cloud/Enrichr/) was used for pathway analysis. TF enrichment analysis was performed using DoRothEA or the ENCODE and ChEA Consensus TFs from ChIP-X in EnrichR. Pathway and TF enrichment values were downloaded as .csv files and plots generated in Prism. GO enrichment analysis to compare cell states was done in R using clusterProlifer^65^ compareClusters function using only statistically significant marker genes.

### Deep Learning to Predict Recovery

In the macrophages and fibroblasts, we exported the log-normalized and scaled counts matrix for the pre-LVAD HF group (U-pre/RR-pre). The counts matrix was filtered to only include *RUNX1* target genes. The target genes were obtained from low and high throughput functional studies (https://maayanlab.cloud/Harmonizome/gene_set/RUNX1/CHEA+Transcription+Factor+Targets). Next, we used the sklearn.model_selection function to split our data into a train and test set (with test.size = 0.3). We then used the StandardScaler function from the sklearn library to scale the train and test feature matrices. We used Keras from tensorflow with a dropout l2 regularization to train a deep neural network to classify nuclei into RR or U category. The model was compiled using a ‘adam’ optimizer and a binary cross-entropy loss function. AUC metrics were calculated. To provide an alternative comparison, we also trained. A random forest classifier using the RandomForestClassifier function from the sklearn.ensemble library. Both models were applied on the test dataset and ROC curves plotted to compare the two approaches. Deep learning was repeated several times and the accuracy metrics were consistent across simulation runs; in each run a different permutation of train/test was used.

### Gene Regulatory Network Construction

CellOracle (v0.10.5)^17^ was used to perform GRN analysis in macrophages and fibroblasts separately. Processed data was imported into scanpy as .h5ad files. The dataset was then down sampled to 20k nuclei to limit computational memory usage. Highly variable genes (3,000) were kept for GRN construction. A promoter DNA sequences base GRN was initialized for humans. Unscaled counts were used to generate the CellOracle object and KNN imputation was performed. Cluster specific GRN was then constructed and the processed GRN was saved for subsequent TF KO analysis.

### In Silico TF Perturbation

For *in silico RUNX1* KO simulation in macrophages and fibroblasts the processed oracle object and inferred GRNs were loaded. The GRN was then fit using ridge regression models for the simulation. To simulate a KO, the *RUNX1* expression was set to 0, Next, we calculated cell state transition probabilities which are visualized as vectors on a digitized grid. To establish a baseline developmental flow field, we use Palantir as described before to calculate pseudotime values. Next, the Gradient_calculator function from the CellOracle library was used to calculate a developmental vector field on the digitized grid. As described in CellOracle, an inner product was calculated between the baseline vector field and the post-KO simulation vector field to assess perturbations in different cell states to understand which populations are enriched and depleted post *RUNX1* KO. All visualization parameters were used as per CellOracle recommendations.

### External In Vivo Dataset Analysis

scRNA-seq analysis: We use publicly available data with annotations from the original manuscript. Specifically, we extracted the fibroblasts (POSTN+ and POSTN-) and myeloid cells. Using the human pseudobulk cardiac recovery genes, we calculated a gene signature for these genes in the mouse dataset and compared sham, TAC, TAC + JQ1, and TAC + JQ1 withdrawn. For the purposes of reference mapping, we extracted the raw filtered matrices for all samples, processed the data using the same pipeline we applied for the human data, and build a single cell map which we annotated using canonical gene markers. This object was then used to annotate the ATAC-seq data.

scATAC-seq analysis: Raw fragment files were used to construct arrow files in ArchR^66^. Briefly, we used ArchR to calculate doublet scores, filtered doublets, only kept cells with TSS > 10, and number of unique fragments > 10,000. Next, we computed an iterative LSI, integrated data with harmony across samples, added clusters, and computed a UMAP embedding. To annotate cells from the ATAC-seq data, we used the addGeneIntegrationMatrix function with the above processed and annotated scRNA-seq reference. Using these cluster annotations, we made pseudobulk replicates with the addGroupCoverages function. Then we used the addReproduciblePeakSet function and macs2 to call peaks. We then subsetted the fibroblasts and used ArchR to compute peak-to-gene links and used browser plots to visualize peak to gene links around the *Runx1* locus. We split the browser track by condition to assess chromatin accessibility with peaks linked to the *Runx1* gene. Within this same locus, we also leveraged publicly available BRD4 and H3K27ac ChIP-seq data^18^ - .bed files were downloaded, and tracts plotted using IGV (v2.13.1).

### Statistics and Reproducibility

Statistical significance was calculated in Prism 9 (Graphpad). For RNAscope image quantification 5 fields were quantified per patient in areas of maximal staining and representative images from those areas shown in the Fig. 3, 4, 5, and 6 and Extended Data Fig. 6, 7, and 8.

## Data Availability

Raw sequencing files can be found on the Gene Expression Omnibus (). Processed data will be made available in a user-friendly format online.

## Code Availability

All scripts used for analysis in this manuscript can be found on GitHub ().

---

**Extended Data Figure 1.**
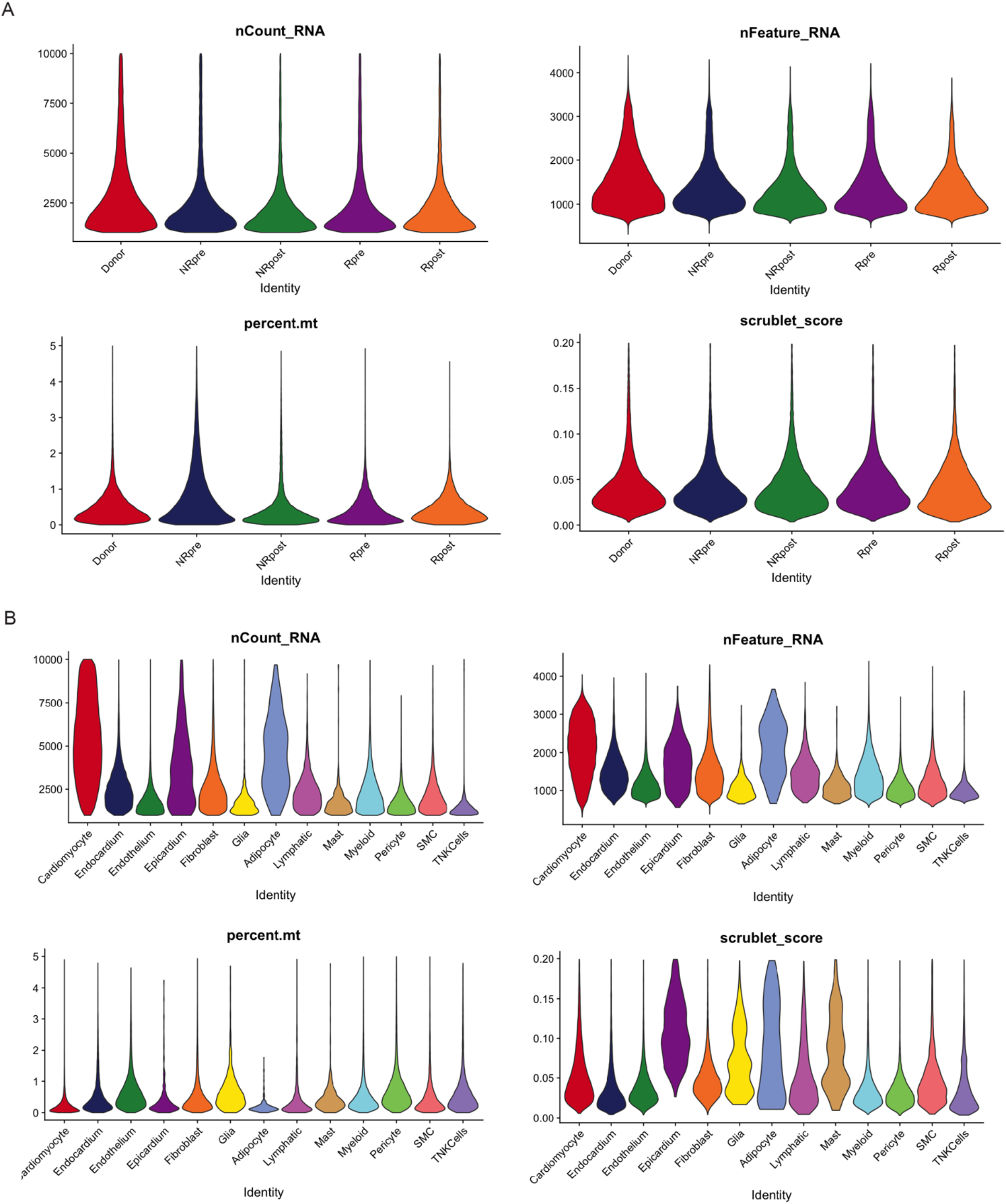
Quality control metrics. nCount_RNA, nFeature_RNA, percent.mt, and scrublet doublet score split by (A) condition and (B) cell type.

**Extended Data Figure 2.**
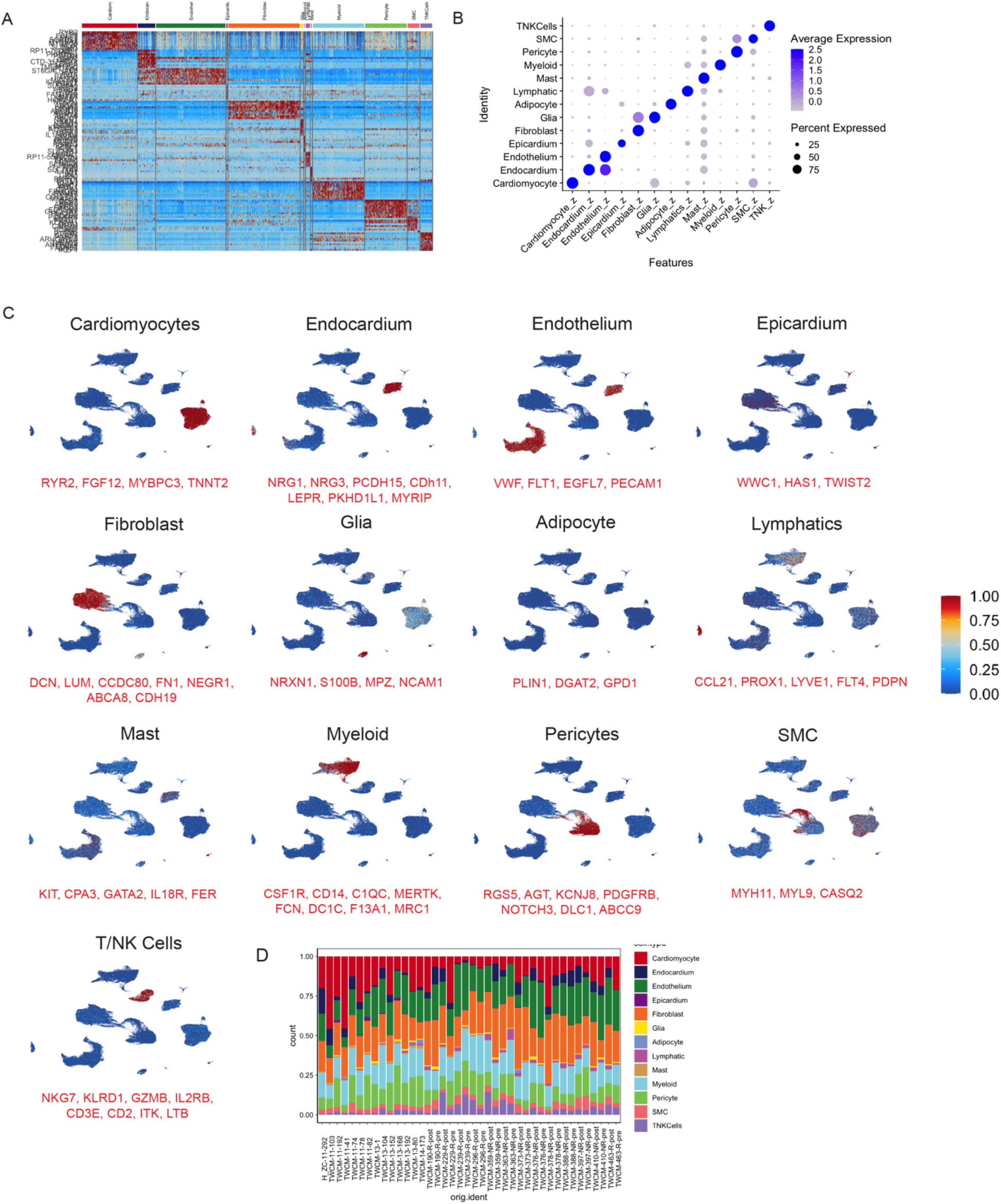
Global clustering. (A) Heatmap of top10 marker gens for each cell type identified via DE analysis. (B) DotPlot for cell type gene set scores from (A) where the x-axis is cell type gene signature and y-axis is the cluster. (C) Gene set z-scores for top gene markers for each cell type plotted in the UMAP embedding. (D) Cell type composition for each of the patient samples.

**Extended Data Figure 3.**
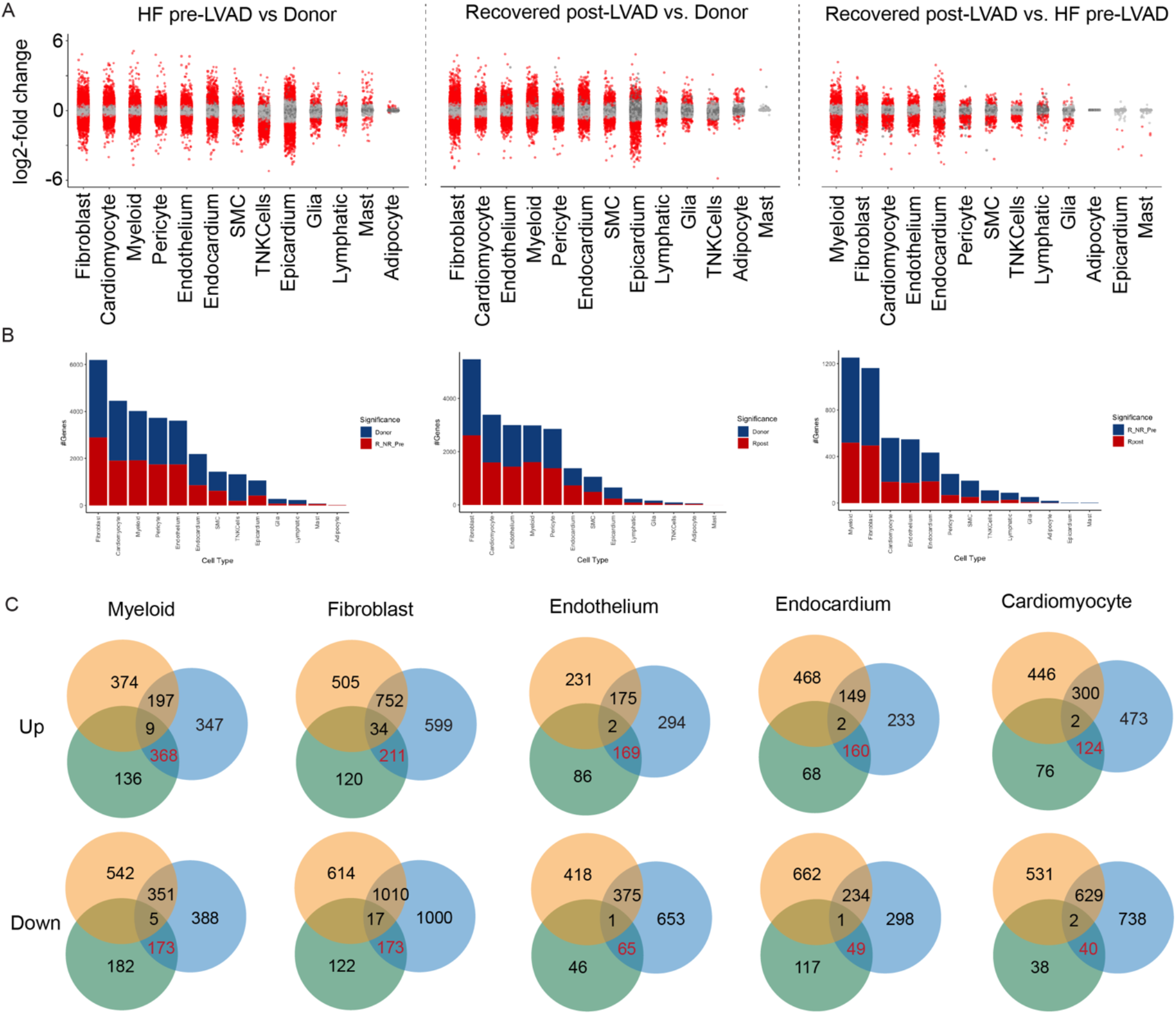
Pseudobulk DE analysis to unravel cardiac recovery. (A) Pseudobulk DE analysis in each cell type in 3 comparison groups: pre-LVAD HF vs donor, RR-post vs donor, and RR-post vs pre-LVAD HF. Red dots indicate statistically significant genes (adjusted p-value < 0.05). (B) Total number of statistically significant (adjusted p-value < 0.05 and log2FC > 0.58) per cell type in comparisons from (A). (C) Number of overlapping genes in five major cell populations which are up and down in the comparisons from (A). Red number is the number of cardiac recovery genes.

**Extended Data Figure 4.**
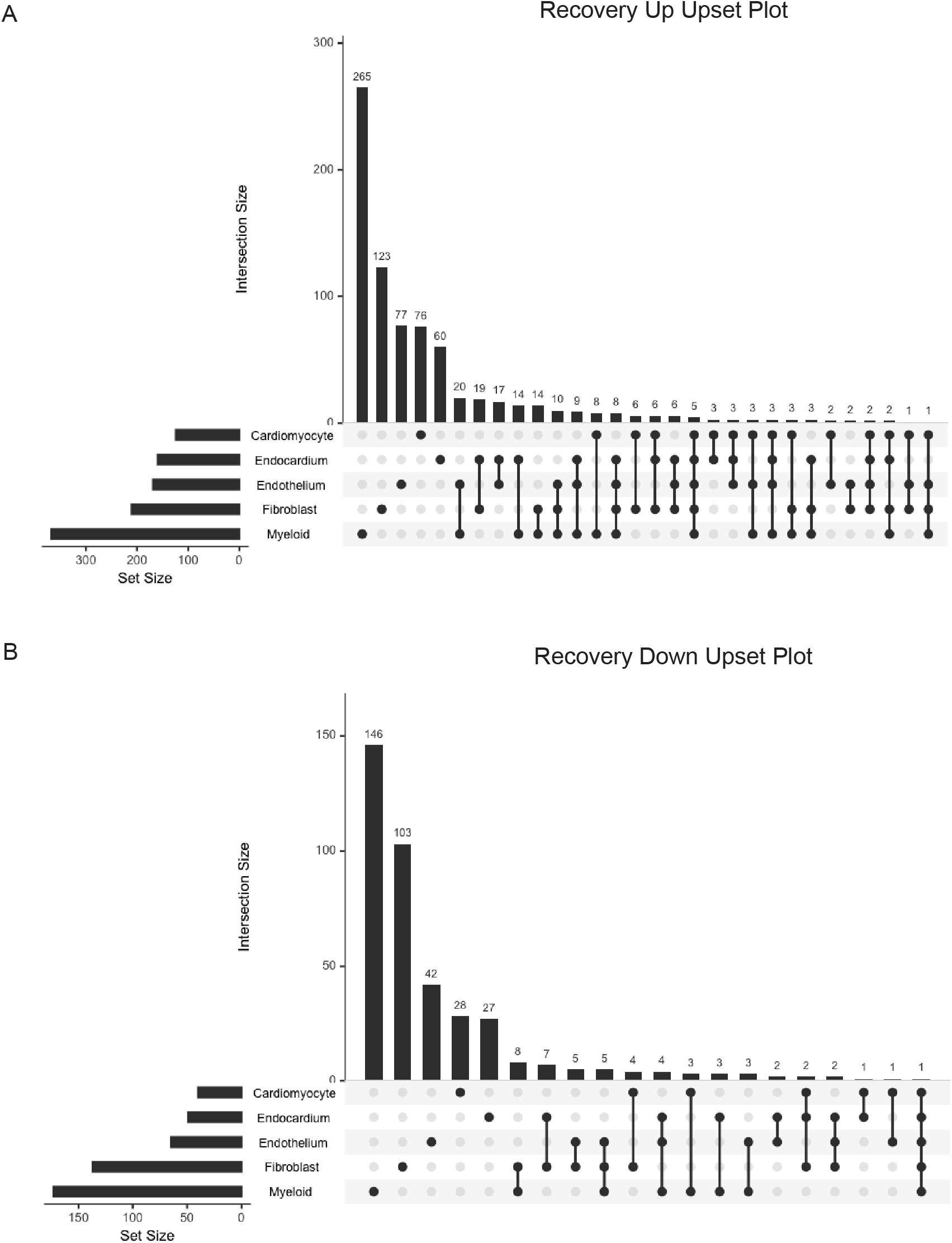
Cardiac recovery overlap amongst cell types. UpSet plot showing overlap in cardiac recovery genes from (Fig.2) in five major cell populations which are (A) up and (B) down in cardiac recovery.

**Extended Data Figure 5.**
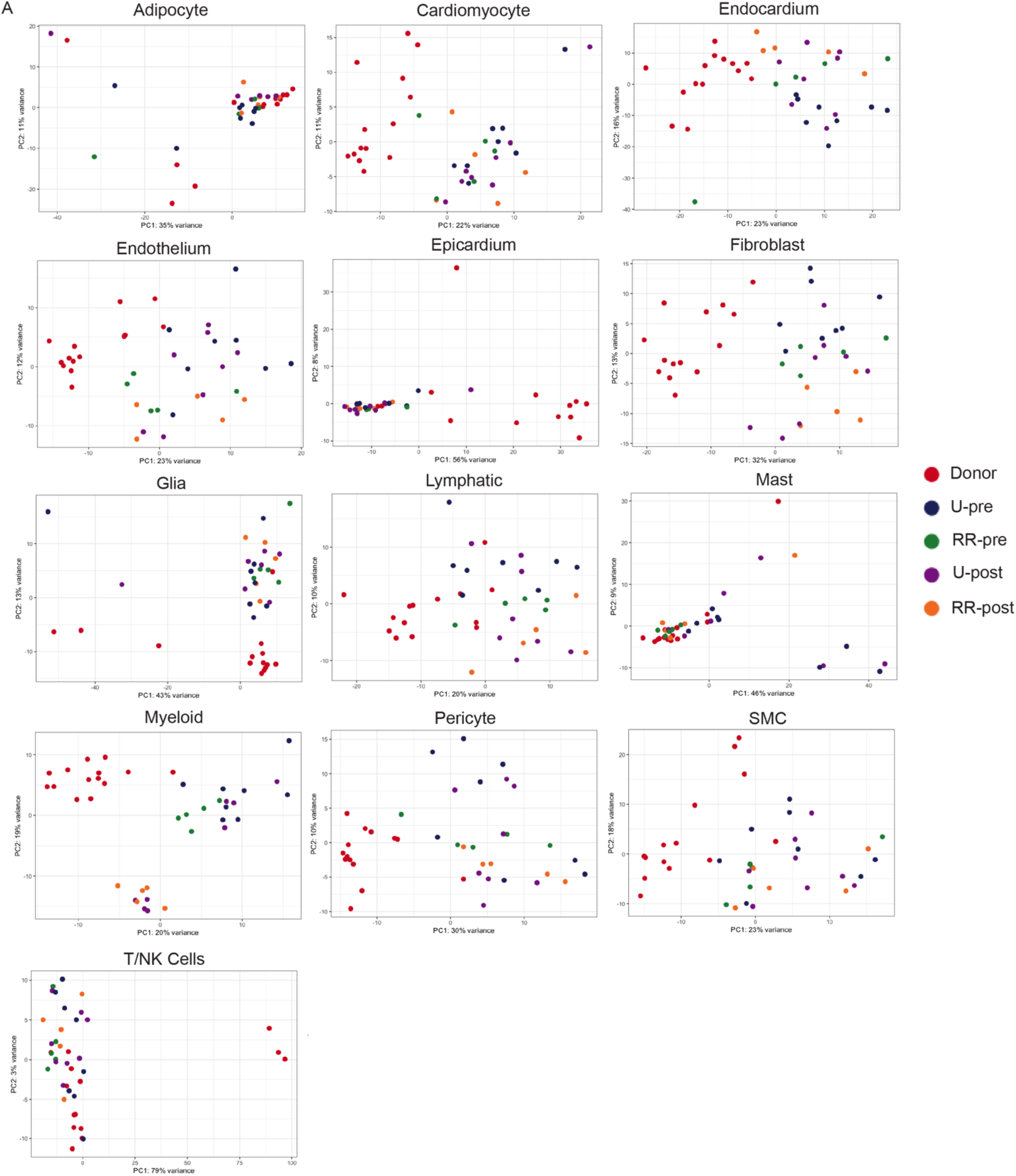
Pseudobulk PCA analysis in each cell type colored by five conditions (donor, U-pre, U-post, RR-pre, and RR-post).

**Extended Data Figure 6.**
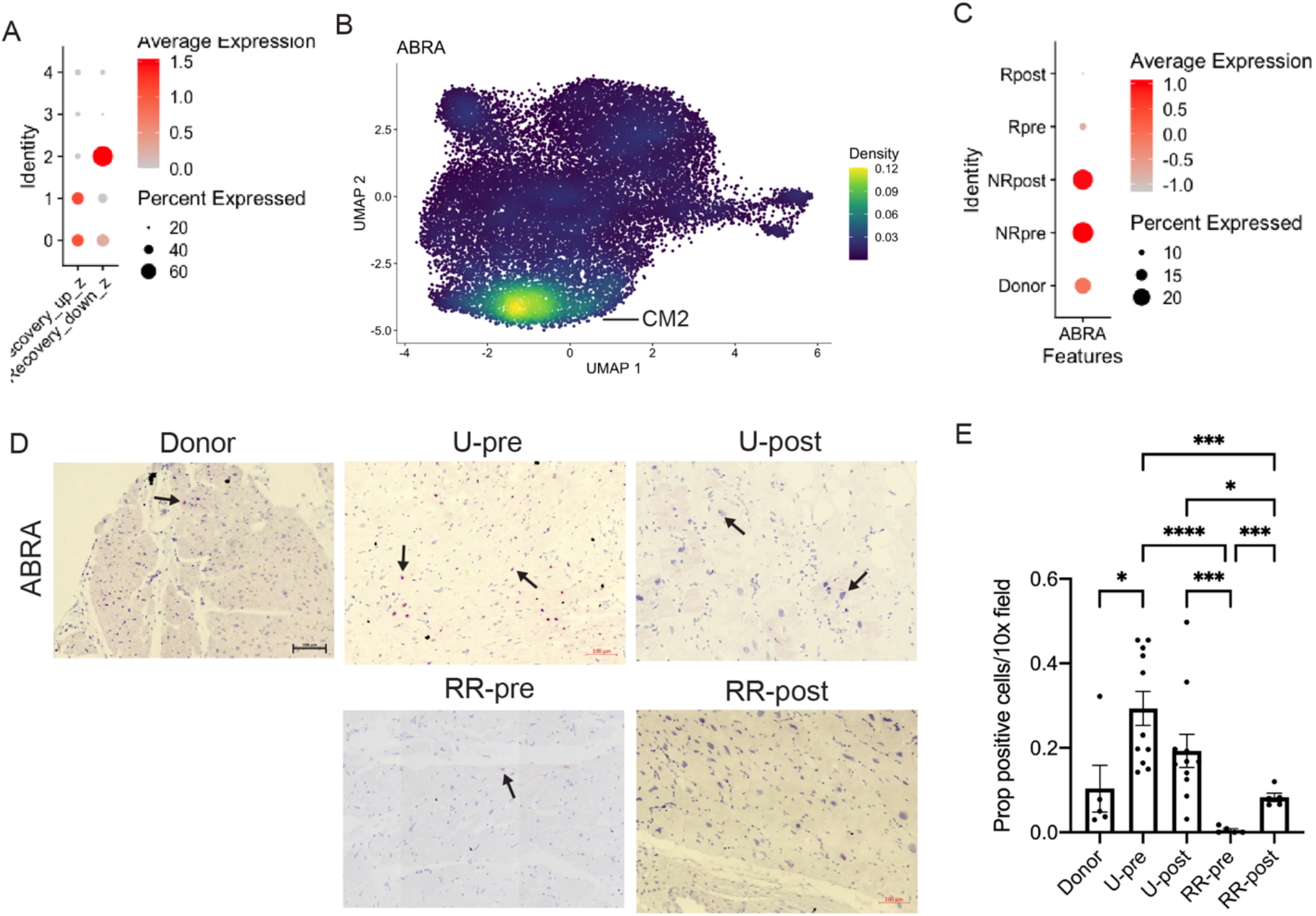
ABRA expression enriched in unloaded group. (A) DotPlot of cardiomyocyte specific recovery up- and down signature grouped by CM cell states. (B) Density plot of ABRA expression in UMAP embedding. (C) DotPlot of ABRA expression in cardiomyocytes grouped by condition. (D) RNAscope images of ABRA in 5 conditions. (E) RNAscope images quantified across an array of patients. Un-paired t-test with Welch’s correction used for comparisons; donor vs U-pre (*P = 0.023), U-pre vs RR-pre (***P < 0.0001), U-pre vs RR-post (***P = 0.0003), U-post vs RR-pre (***P = 0.0007), U-post vs RR-post (*P = 0.0188), and RR-pre vs RR-post (***P = 0.0007).

**Extended Data Figure 7.**
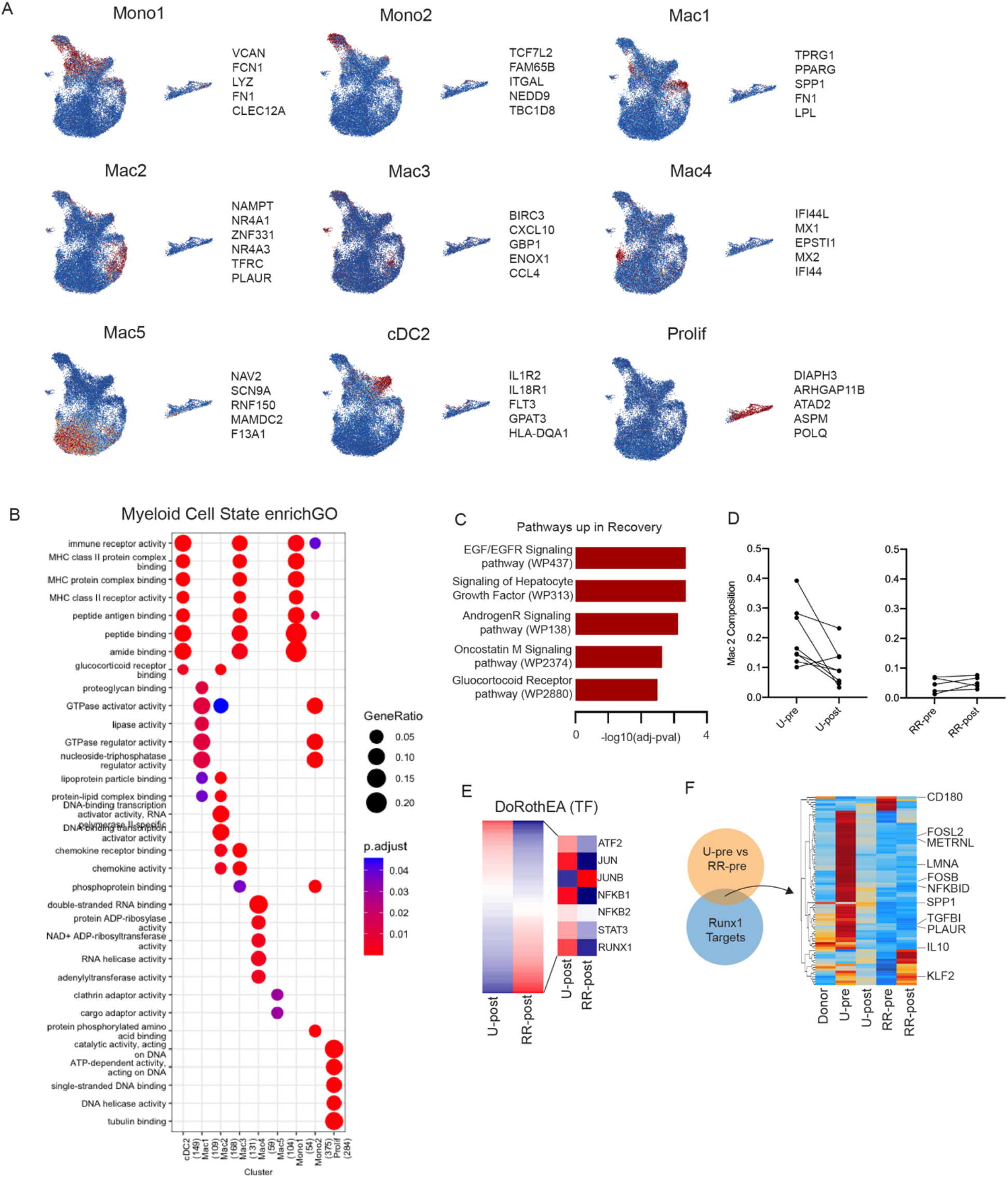
Macrophage diversity in recovery. (A) Gene set z-scores for top gene markers for each cell state plotted in the UMAP embedding. (B) Enrich GO using compareclusters from cluster Prolifer across macrophage cell states. (C) WikiPathways enriched in cardiac recovery. (D) Paired comparison of Mac 2 cluster composition at patient level split by U and RR group. (E) DoRothEA TF enrichment analysis in U-post and RR-post zoomed in on some key differentially enriched TFs. (F) Overlap between Runx1 target genes and DE genes between U-pre and RR-pre with heatmap of respective genes split by condition.

**Extended Data Figure 8.**
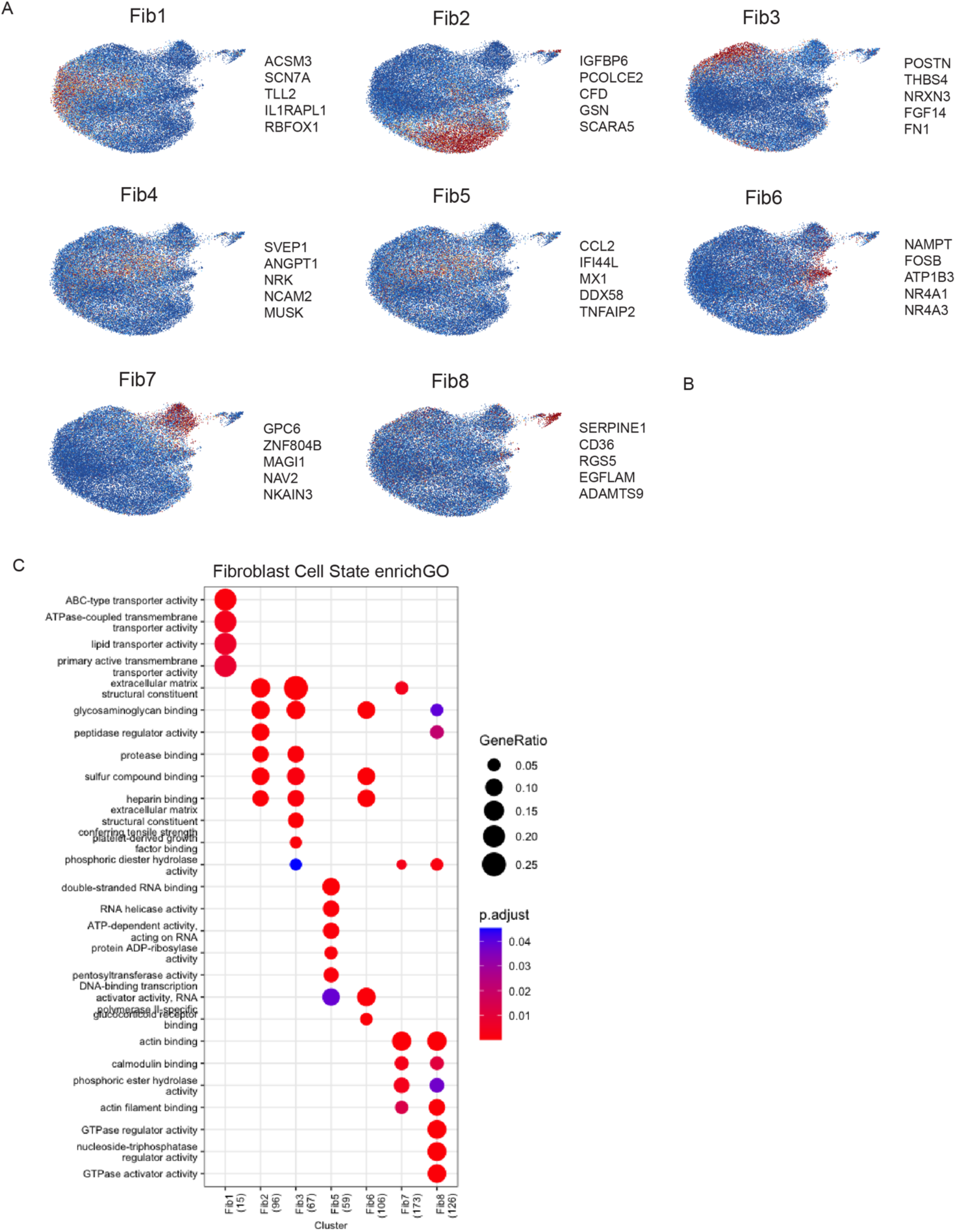
Fibroblast diversity in recovery. (A) Gene set z-scores for top gene markers for each cell state plotted in the UMAP embedding. (B) DotPlot for cell type gene set scores from (A) where the x-axis is cell type gene signature and y-axis is the cluster. (C) Enrich GO using compareclusters from cluster Prolifer across fibroblast cell states.

**Extended Data Figure 9.**
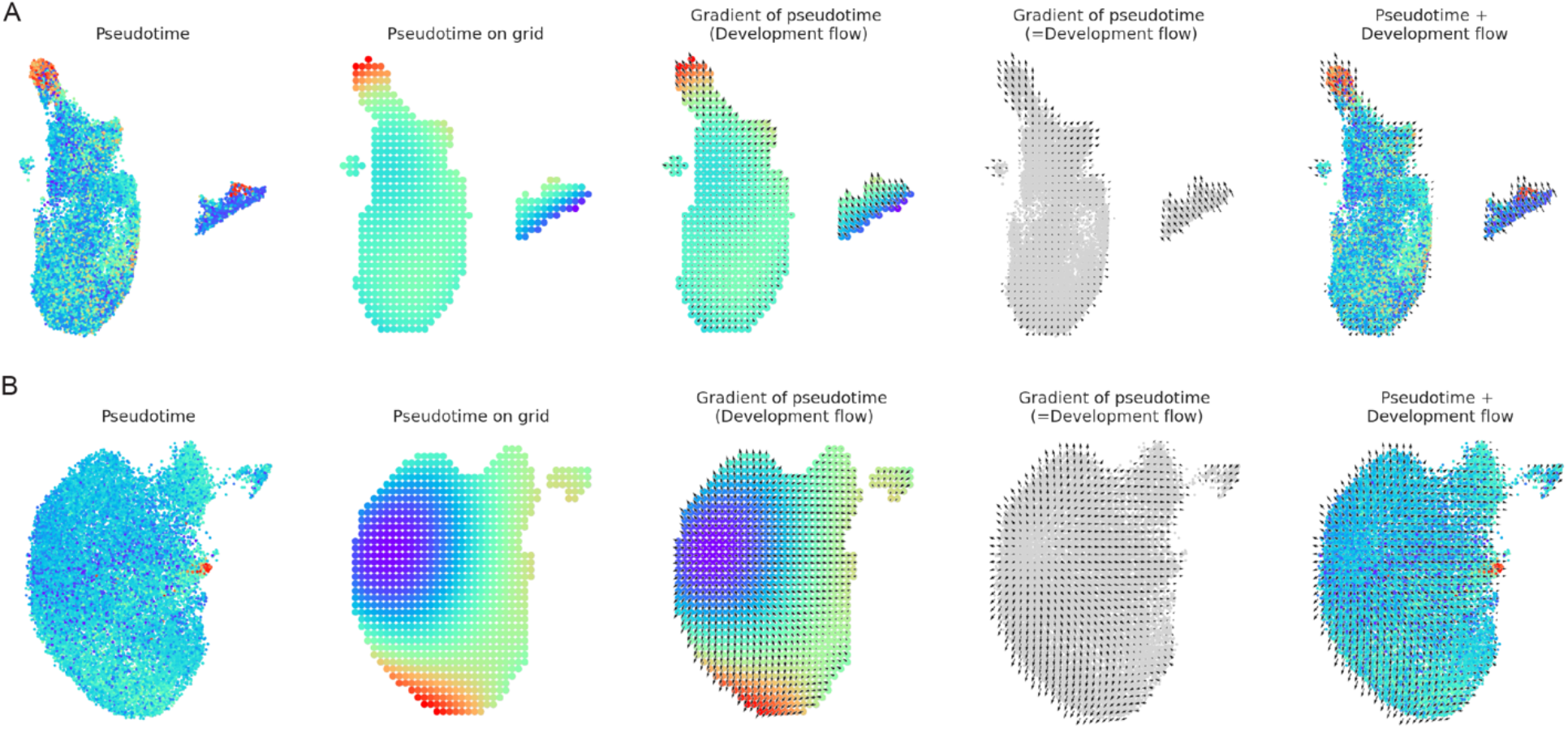
CellOracle developmental flow and digitized grid construction in (A) Macrophages and (B) Fibroblasts.

## References

1. Roger, V. L. Epidemiology of Heart Failure. Circ. Res. 113, 646 (2013).

2. Jessup, M. & Brozena, S. Heart Failure. https://doi.org/10.1056/NEJMra021498 348, 2007–2018 (2003).

3. Topkara, V. K. et al. Myocardial Recovery in Patients Receiving Contemporary Left Ventricular Assist Devices: Results from the Interagency Registry for Mechanically Assisted Circulatory Support (INTERMACS). Circ. Heart Fail. 9, (2016).

4. Burkhoff, D., Topkara, V. K., Sayer, G. & Uriel, N. Reverse Remodeling With Left Ventricular Assist Devices. Circ. Res. 128, 1594–1612 (2021).

5. Selzman, C. H. et al. Bridge to Removal: A Paradigm Shift for Left Ventricular Assist Device Therapy. Ann. Thorac. Surg. 99, 360 (2015).

6. Givertz, M. M. & Mann, D. L. Epidemiology and natural history of recovery of left ventricular function in recent onset dilated cardiomyopathies. Curr. Heart Fail. Rep. 10, 321–330 (2013).

7. Kanwar, M. K. et al. Clinical myocardial recovery in advanced heart failure with long term left ventricular assist device support. J. Hear. Lung Transplant. (2022). doi:10.1016/J.HEALUN.2022.05.015

8. Shepherd, C. W. & While, A. E. Cardiac rehabilitation and quality of life: a systematic review. Int. J. Nurs. Stud. 49, 755–771 (2012).

9. Tseliou, E. et al. Biology of myocardial recovery in advanced heart failure with long-term mechanical support. J. Hear. Lung Transplant. (2022). doi:10.1016/J.HEALUN.2022.07.007

10. McGuire, A. L. et al. The road ahead in genetics and genomics. Nat. Rev. Genet. 2020 2110 21, 581–596 (2020).

11. Eraslan, G. et al. Single-nucleus cross-tissue molecular reference maps toward understanding disease gene function. Science (80-.). 376, (2022).

12. Litviňuková, M. et al. Cells of the adult human heart. Nat. 2020 5887838 588, 466–472 (2020).

13. Koenig, A. L. et al. Single-cell transcriptomics reveals cell-type-specific diversification in human heart failure. Nat. Cardiovasc. Res. 2022 13 **1**, 263–280 (2022).

14. Chaffin, M. et al. Single-nucleus profiling of human dilated and hypertrophic cardiomyopathy. Nature (2022). doi:10.1038/S41586-022-04817-8

15. Tucker, N. R. et al. Transcriptional and Cellular Diversity of the Human Heart. Circulation 142, 466–482 (2020).

16. Nicin, L. et al. A human cell atlas of the pressure-induced hypertrophic heart. Nat. Cardiovasc. Res. 2022 12 **1**, 174–185 (2022).

17. Kamimoto, K. et al. Gene Regulatory Network Reconfiguration in Direct Lineage Reprogramming. bioRxiv 2022.07.01.497374 (2022). doi:10.1101/2022.07.01.497374

18. Alexanian, M. et al. A transcriptional switch governs fibroblast activation in heart disease. Nat. 2021 5957867 595, 438–443 (2021).

19. Gupta, D. K. et al. Assessment of myocardial viability and left ventricular function in patients supported by a left ventricular assist device. J. Heart Lung Transplant. 33, 372– 381 (2014).

20. Burke, M. A. & Givertz, M. M. Assessment and management of heart failure after left ventricular assist device implantation. Circulation 129, 1161–1166 (2014).

21. Bajpai, G. et al. Tissue Resident CCR2- and CCR2+ Cardiac Macrophages Differentially Orchestrate Monocyte Recruitment and Fate Specification Following Myocardial Injury. Circ. Res. 124, 263–278 (2019).

22. Zaman, R. et al. Selective loss of resident macrophage-derived insulin-like growth factor-1 abolishes adaptive cardiac growth to stress. Immunity 54, 2057–2071.e6 (2021).

23. Dick, S. A. et al. Self-renewing resident cardiac macrophages limit adverse remodeling following myocardial infarction. Nat. Immunol. 20, 29 (2019).

24. Garcia-Alonso, L., Holland, C. H., Ibrahim, M. M., Turei, D. & Saez-Rodriguez, J. Benchmark and integration of resources for the estimation of human transcription factor activities. Genome Res. 29, 1363–1375 (2019).

25. Asleh, R., Amir, O. & Kushwaha, S. S. Dynamics of myocardial fibrosis after left ventricular assist device implantation: should speeding up the scar have us scared stiff? Eur. J. Heart Fail. 23, 335–338 (2021).

26. Wilcox, J. E. et al. “Targeting the Heart” in Heart Failure: Myocardial Recovery in Heart Failure With Reduced Ejection Fraction. JACC Hear. Fail. 3, 661–669 (2015).

27. Stratton, M. S. et al. Dynamic Chromatin Targeting of BRD4 Stimulates Cardiac Fibroblast Activation. Circ. Res. 125, 662 (2019).

28. Aghajanian, H. et al. Targeting cardiac fibrosis with engineered T cells. Nat. 2019 5737774 573, 430–433 (2019).

29. Bocchi, V. D. et al. The coding and long noncoding single-cell atlas of the developing human fetal striatum. Science (80-.). 372, (2021).

30. Rose, E. A. et al. Long-Term Use of a Left Ventricular Assist Device for End-Stage Heart Failure. https://doi.org/10.1056/NEJMoa012175 345, 1435–1443 (2001).

31. Miller, L., Birks, E., Guglin, M., Lamba, H. & Frazier, O. H. Use of Ventricular Assist Devices and Heart Transplantation for Advanced Heart Failure. Circ. Res. 124, 1658– 1678 (2019).

32. Dharmavaram, N. et al. National trends in heart donor usage rates: Are we efficiently transplanting more hearts? J. Am. Heart Assoc. 10, 19655 (2021).

33. Bowen, R. E. S., Graetz, T. J., Emmert, D. A. & Avidan, M. S. Statistics of heart failure and mechanical circulatory support in 2020. Ann. Transl. Med. 8, 827–827 (2020).

34. Drakos, S. G. et al. Bridge to Recovery: Understanding the Disconnect Between Clinical and Biological Outcomes. Circulation 126, 230 (2012).

35. Lenneman, A. J. & Birks, E. J. Treatment strategies for myocardial recovery in heart failure. Curr. Treat. Options Cardiovasc. Med. 16, (2014).

36. Halliday, B. P. et al. Withdrawal of pharmacological treatment for heart failure in patients with recovered dilated cardiomyopathy (TRED-HF): an open-label, pilot, randomised trial. Lancet 393, 61–73 (2019).

37. Bottle, A., Faitna, P., Aylin, P. P. & Cowie, M. R. Original research: Five-year outcomes following left ventricular assist device implantation in England. Open Hear. 8, 1658 (2021).

38. Wang, L. et al. Single-cell reconstruction of the adult human heart during heart failure and recovery reveals the cellular landscape underlying cardiac function. Nat. Cell Biol. 2020 221 **22**, 108–119 (2020).

39. Birks, E. J. et al. Prospective Multicenter Study of Myocardial Recovery Using Left Ventricular Assist Devices (RESTAGE-HF [Remission from Stage D Heart Failure]) Medium-Term and Primary End Point Results. Circulation 142, 2016–2028 (2020).

40. Zhang, J. & Narula, J. Molecular biology of myocardial recovery. Surg. Clin. North Am. 84, 223–242 (2004).

41. Klotz, S., Jan Danser, A. H. & Burkhoff, D. Impact of left ventricular assist device (LVAD) support on the cardiac reverse remodeling process. Prog. Biophys. Mol. Biol. 97, 479– 496 (2008).

42. Wohlschlaeger, J. et al. Reverse remodeling following insertion of left ventricular assist devices (LVAD): A review of the morphological and molecular changes. Cardiovasc. Res. 68, 376–386 (2005).

43. Weinheimer, C. J. et al. Load-Dependent Changes in Left Ventricular Structure and Function in a Pathophysiologically Relevant Murine Model of Reversible Heart Failure. Circ. Heart Fail. 11, e004351 (2018).

44. Bajpai, G. et al. The human heart contains distinct macrophage subsets with divergent origins and functions. Nat. Med. 24, 1234–1245 (2018).

45. Epelman, S., Lavine, K. J. & Randolph, G. J. Origin and Functions of Tissue Macrophages. Immunity 41, 21–35 (2014).

46. Lavine, K. J. et al. Distinct macrophage lineages contribute to disparate patterns of cardiac recovery and remodeling in the neonatal and adult heart. Proc. Natl. Acad. Sci. U. S. A. 111, 16029–16034 (2014).

47. Wong, N. R. et al. Resident cardiac macrophages mediate adaptive myocardial remodeling. Immunity 54, 2072–2088.e7 (2021).

48. Kuppe, C. et al. Decoding myofibroblast origins in human kidney fibrosis. Nature 589, 281–286 (2021).

49. Khalil, H. et al. Fibroblast-specific TGF-β–Smad2/3 signaling underlies cardiac fibrosis. J. Clin. Invest. 127, 3770–3783 (2017).

50. Medzhitov, R. Origin and physiological roles of inflammation. Nature 454, 428–435 (2008).

51. Tzahor, E. & Dimmeler, S. A coalition to heal—the impact of the cardiac microenvironment. Science (80-.). 377, (2022).

52. Sood, R., Kamikubo, Y. & Liu, P. Role of RUNX1 in hematological malignancies. Blood 129, 2070 (2017).

53. Chen, M. J., Yokomizo, T., Zeigler, B. M., Dzierzak, E. & Speck, N. A. Runx1 is required for the endothelial to haematopoietic cell transition but not thereafter. Nat. 2009 4577231 457, 887–891 (2009).

54. Koth, J. et al. Runx1 promotes scar deposition and inhibits myocardial proliferation and survival during zebrafish heart regeneration. Development 147, (2020).

55. Hu, B. et al. Origin and function of activated fibroblast states during zebrafish heart regeneration. Nat. Genet. 2022 1–11 (2022). doi:10.1038/s41588-022-01129-5

56. Amrute, J. M. et al. Cell specific peripheral immune responses predict survival in critical COVID-19 patients. Nat. Commun. 2022 131 **13**, 1–11 (2022).

57. Kong, Y. & Yu, T. A Deep Neural Network Model using Random Forest to Extract Feature Representation for Gene Expression Data Classification. Sci. Rep. (2018). doi:10.1038/s41598-018-34833-6

58. Khouri-Farah, N., Guo, Q., Morgan, K., Shin, J. & Li, J. Y. H. Integrated single-cell transcriptomic and epigenetic study of cell state transition and lineage commitment in embryonic mouse cerebellum. Sci. Adv. 8, (2022).

59. Butler, A., Hoffman, P., Smibert, P., Papalexi, E. & Satija, R. Integrating single-cell transcriptomic data across different conditions, technologies, and species. Nat. Biotechnol. 36, 411 (2018).

60. Hafemeister, C. & Satija, R. Normalization and variance stabilization of single-cell RNA-seq data using regularized negative binomial regression. Genome Biol. 20, 1–15 (2019).

61. Korsunsky, I. et al. Fast, sensitive and accurate integration of single-cell data with Harmony. Nat. Methods 2019 1612 **16**, 1289–1296 (2019).

62. Love, M. I., Huber, W. & Anders, S. Moderated estimation of fold change and dispersion for RNA-seq data with DESeq2. Genome Biol. 15, 1–21 (2014).

63. Wang, F. et al. RNAscope: A Novel in Situ RNA Analysis Platform for Formalin-Fixed, Paraffin-Embedded Tissues. J. Mol. Diagn. 14, 22 (2012).

64. Setty, M. et al. Characterization of cell fate probabilities in single-cell data with Palantir. Nat. Biotechnol. 2019 374 **37**, 451–460 (2019).

65. Wu, T. et al. clusterProfiler 4.0: A universal enrichment tool for interpreting omics data. Innov. 2, 100141 (2021).

66. Granja, J. M. et al. ArchR is a scalable software package for integrative single-cell chromatin accessibility analysis. Nat. Genet. 2021 533 **53**, 403–411 (2021).

